# Kinetic Characterization and Computational Modeling of the *Escherichia coli* Heptosyltransferase II: Exploring the Role of Protein Dynamics in Catalysis for a GT-B Glycosyltransferase

**DOI:** 10.1101/2022.06.13.495986

**Authors:** Bakar A. Hassan, Zhiqi A. Liu, Jozafina Milicaj, Mia S. Kim, Meka Tyson, Yuk Y. Sham, Erika A. Taylor

## Abstract

Glycosyltransferases (GTs) are enzymes that are uniquely adapted to promote the formation of a glycosidic bond between a sugar molecule and a wide variety of substrates. Heptosyltransferase II (HepII) is a GT involved in the lipopolysaccharide (LPS) biosynthetic pathway that transfers the seven-carbon sugar (L-*glycero*-D-*manno*-heptose; Hep) onto a lipid anchored glycopolymer (heptosylated Kdo_2_-Lipid A, Hep-Kdo_2_-Lipid A or HLA). LPS plays a key role in Gram-negative bacterial sepsis as a stimulator of the human immune response and has been used as an adjuvant in vaccines. As such, ongoing efforts towards inhibition of LPS biosynthetic enzymes to aid development of novel antimicrobial therapeutics has driven significant effort towards the characterization of these enzymes. Three heptosyltransferases are involved in the inner-core biosynthesis, with *E. coli* HepII being the last to be quantitatively characterized *in vivo*, as described herein. HepII shares modest sequence similarity with heptosyltransferase I (HepI) while maintaining a high degree of structural homology. Here we report the first kinetic and biophysical characterization of HepII and demonstrate the properties of HepII that are shared by HepI to include sugar donor promiscuity, and sugar acceptor induced secondary structural changes which results in significant thermal stabilization. HepII also has an increased catalytic efficiency and a significantly tighter binding affinity for both of its substrates, with an insensitivity to the number of acyl chains on the sugar acceptor. Additionally, a structural model of the HepII ternary complex, refined by molecular dynamics simulations, was developed to probe potentially important substrate-protein contacts and revealed the potential of Tryptophan (Trp) residues responsible for reporting on ligand binding. As was previously described for HepI, Tryptophan fluorescence in HepII allowed observation of substrate induced changes in Trp fluorescence intensity which enabled determination of substrate dissociation constants. Combined, these efforts meaningfully enhance our understanding of the Heptosyltransferase family of enzymes and will aid in future efforts to design novel, potent and specific inhibitors for this family of enzymes.

## Introduction

The ever growing concern of multi-drug resistant bacterial infections is invigorating efforts towards the development of novel antimicrobials that could be used as standalone treatments or as part of combinatorial therapeutics.^1–3^ In Gram-negative bacteria, the lipopolysaccharide (LPS) is a major contributor to virulence and it acts as a barrier to protect against xenobiotics including hydrophobic antibiotics.^4^ The LPS is a lipid anchored glycopolymer that is incorporated into the outer leaflet of the outer membrane of Gram-negative bacteria. LPS can elicit the human immune response through interactions with toll-like receptor 4 (TLR4) and plays a role in bacterial sepsis.^5^ This modulatory behavior has inspired some to use LPS and various analogs as vaccine adjuvants.^6^ The fully synthesized LPS is a major component of the outer leaflet of the outer membrane, making up approximately 75% of the bacterial membrane surface,^17^ with LPS having been shown to be important for the formation of bacterial biofilms in multiple organisms,^7^ making the LPS biosynthetic enzymes targets for inhibitor development to enable multiple medical applications.^8–12^ The LPS consist of the acylated Lipid A base, an inner core and outer core oligosaccharide region, and O-antigen repeating region (Figure 1A). The LPS Lipid A and core region are both assembled on the inner leaflet of the inner membrane, while the O-antigen is added after the truncated LPS is transported onto the bacteria’s surface via cross membrane transporters (Figure 1B).^15–16^ The Lipid A and inner core are highly conserved among Gram-negative bacteria making them good targets for the design of potential LPS biosynthesis inhibitors that would have broad spectrum activity. Previous *in vivo* experiments have demonstrated that mutations that lead to truncation of LPS at the inner core exhibit a phenotype that includes an increased susceptibility to hydrophobic antibiotics due to a compromised outer membrane.^18^ Heptosyltransferase I and II (HepI, HepII), enzymes involved in the inner core biosynthesis, are found in a wide genus of clinically relevant bacteria including Enterobacteriaceae, Pasteurellaceae, Pseudomonadaceae, Moraxellaceae, Vibrionaceae, Burkholderiaceae and Neisseriaceae.^19^ Therefore, inhibition of HepI or HepII *in vivo* would lead to a truncated LPS^20^ making these enzymes highly desirable drug targets.

**Figure 1.**
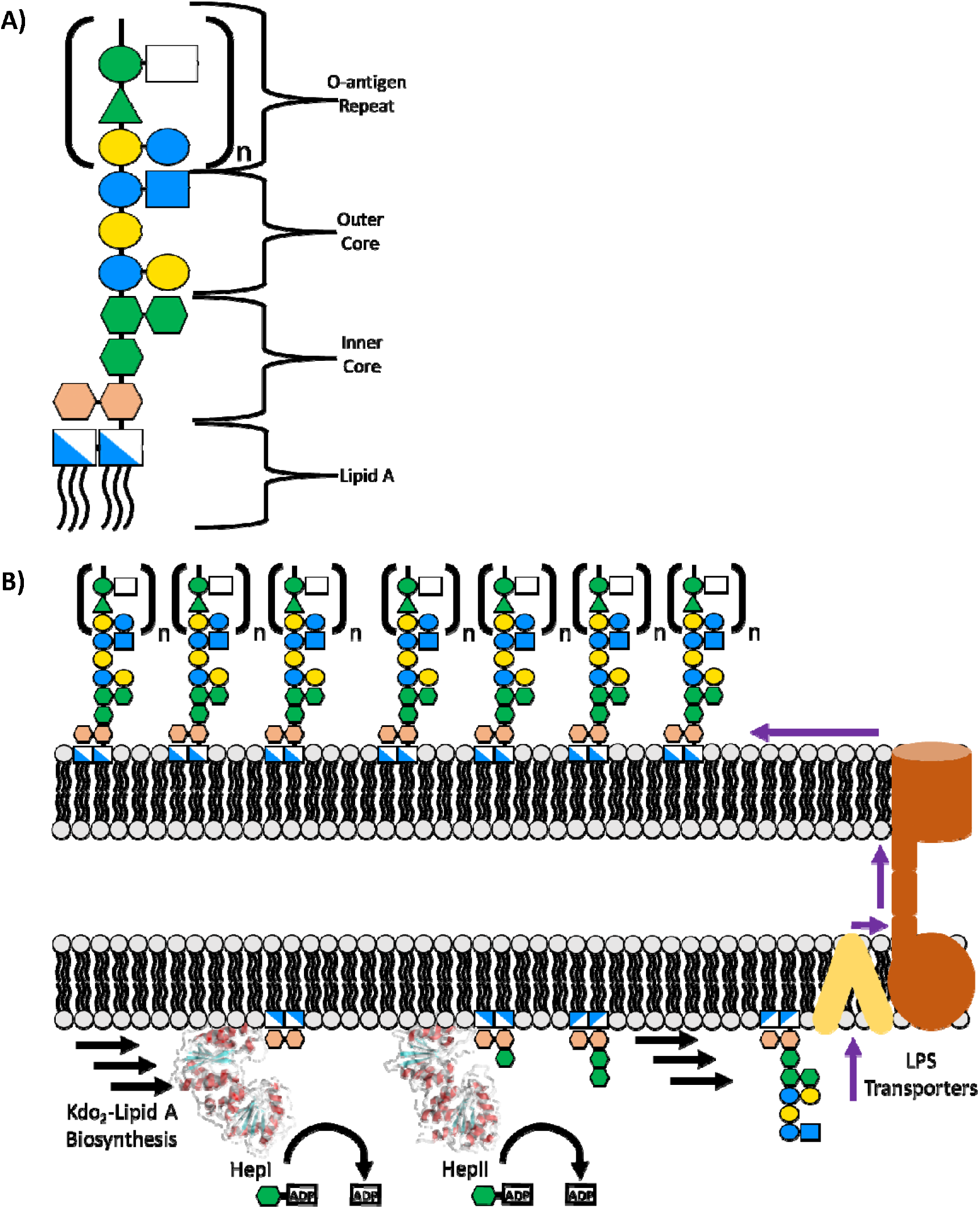
Lipopolysaccharide components and inner core biosynthesis. **(A)** Regions of the full lipopolysaccharide (LPS) and their constituent saccharides. **(B)** Cartoon representation of the role of Heptosyltransferase I and II and their association with the inner membrane of the intracellular side to catalyze the transfer of heptose moiety onto the elongating glycopolymer (bottom left) prior to transportation (bottom/top right) and embedding of the LPS into the outer membrane (top left).

HepI has been thoroughly characterized both *in vivo* and *in vitro*, with some nanomolar inhibitors successfully designed for it to date.^21–31^ HepII, however, had previously only been functionally characterized *in vivo*, with an apo crystal structure (PDB: 1PSW; Uniprot: P37692) with no corresponding manuscript.^26,32–33^ HepI and HepII function consecutively in the LPS inner core biosynthetic pathway. HepI utilizes an ADP-L-*glycero*-β-D-*manno*-heptose (ADP-Hep) and transfers the heptose moiety onto the Kdo_2_-Lipid A forming an α(1->5) glycosidic bond yielding ADP and Hep-Kdo_2_-Lipid A (Figure 1B). HepII utilizes an identical sugar donor ADP-Hep and transfers the heptose moiety onto the Hep-Kdo_2_-Lipid A (product of HepI reaction) forming an α(1->3) glycosidic bond to form ADP and Hep_2_-Kdo_2_-Lipid A (Figure 1-2). These enzymes catalyze near identical reactions while having a surprisingly low degree of sequence identity (32%; Figure S1). They are both part of the GT-B structural class of glycosyltransferases and the GT-9 structural family in the Carbohydrate-Active enZYme (CaZY) database (www.cazy.org). As members of the GT-B structural class, they have two independent α/β/α Rossmann-like domains connected by a linker region (Figure 3). As in all GT-B enzymes, the N-terminal domain binds the nucleophilic, sugar acceptor substrate and the C-terminal domain binds the activated sugar donor. It was previously demonstrated that HepI can utilize a hexose analog sugar donor, ADP-β-mannose (ADP-Man),^31^ while also tolerating removal of acyl chains from the acceptor;^24^ these two prior results motivated our investigation of the substrate selectivity of HepII. Additionally, qualitative *in vivo* data has demonstrated HepII’s capabilities of utilizing ADP-Man as an alternative sugar donor while also demonstrating that the enzyme has a greater catalytic efficiency relative to HepI; however, these findings have never before been validated *in vitro*.^26^ Herein, we report the first characterization of *Escherichia coli* HepII including kinetic and fluorescence based *in vitro* studies, as well as computational modeling and molecular dynamics *in silico* methods to gain a greater insight into the structural properties that govern the functional characteristics of this enzyme. The study of HepII will enable the future design of inhibitors that act as potent and specific antimicrobials with synergistic properties in combinatorial therapeutics with other hydrophobic antibiotics.

**Figure 2.**
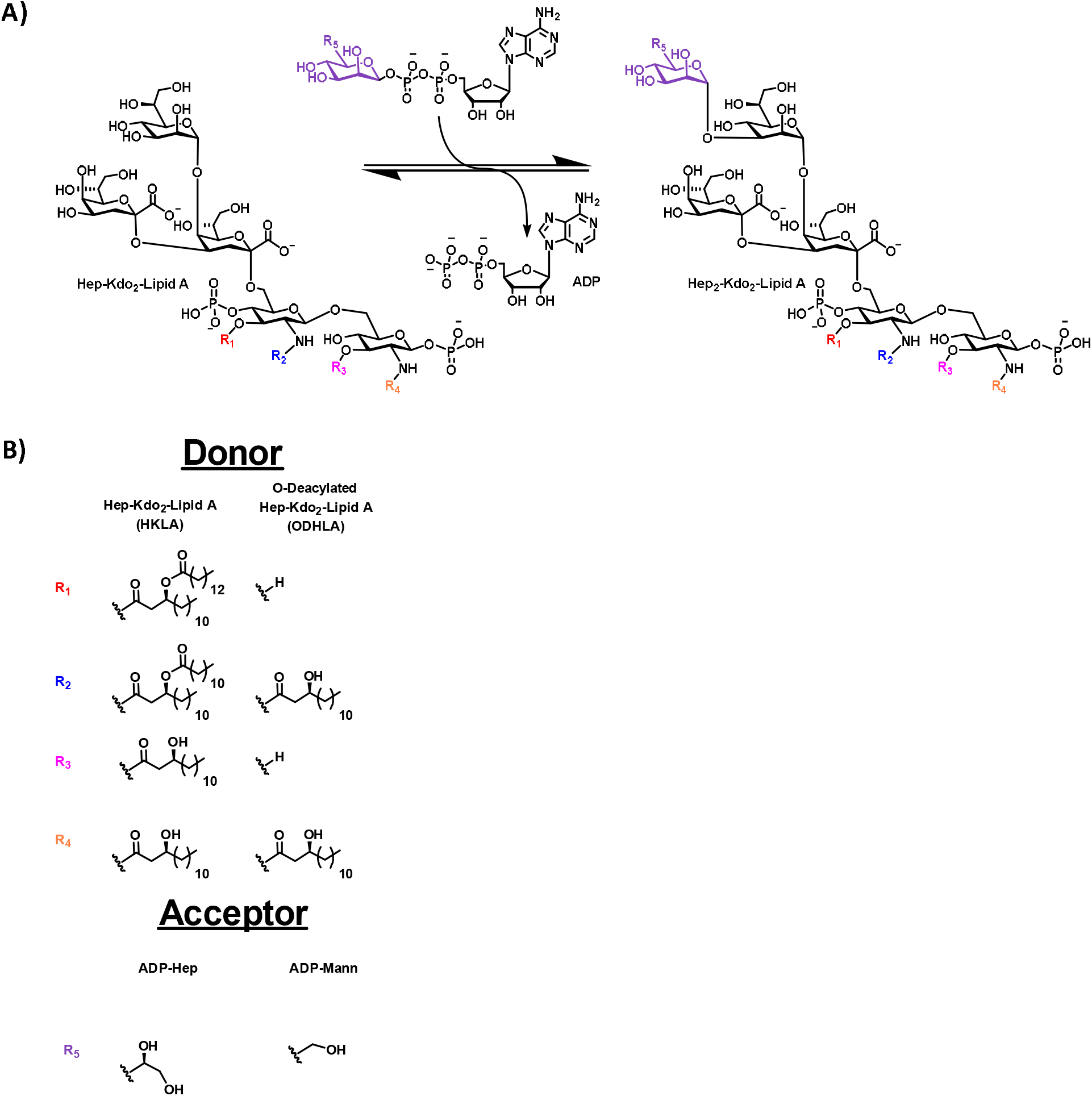
Reaction catalyzed by HepII with various substrates. **(A)** Reaction catalyzed by Heptosyltransferase II where a 6 or 7 carbon saccharide is transferred onto a Hep-Kdo_2_-Lipid A with the release of ADP to form the product, Hep_2_-Kdo_2_-Lipid A. **(B)** The Hep-Kdo_2_-Lipid A can be deacylated at the O positions (R1 and R3) to aid with solubility and both the 7 carbon native donor and a 6 carbon analogue were tested (R5).

**Figure 3.**
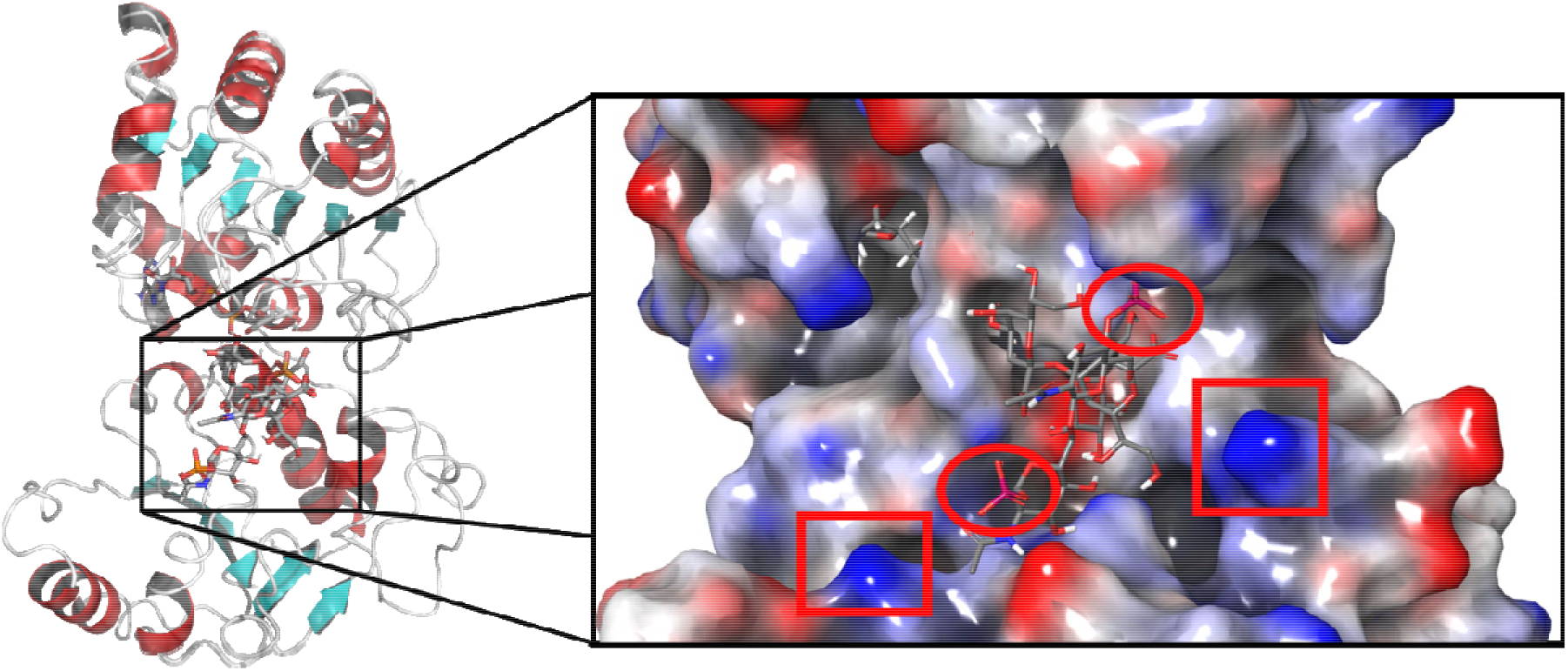
Hybrid ternary complex model. A ternary complex model of HepII was generated based on the crystal structure of the apo (PDB: 1PSW) with both ligands modeled in their respective domains (left). An electrostatic potential surface of HepII that demonstrates the positively charged surface of the active site (boxed lysine/arginine residues) to accommodate binding of the acceptor (circled phosphate groups of Hep-Kdo2-Lipid A) (right).

## Materials and Methods

### Multiple Sequence Alignment and Sequence Conservation Analysis

Multiple Sequence alignments and sequence conservation of HepI and HepII were obtained from the consurf webserver.^34–37^ Briefly, *E. coli* HepI and HepII amino acid sequences were obtained from crystal structures (HepI: PDB 6DFE, HepII: PDB 1PSW). Homologous sequences for each protein were retrieved from the uniref90 server and yielded 2028 unique sequences for HepI and 1994 unique sequences for HepII using a sequence similarity cutoff including only sequences with 95% to 35% sequence identity. Sequences were ordered by ascending E values and 150 representative sequences were chosen equally spaced from the list of available sequences. Multiple sequence alignment for homologues and pairwise sequence alignment between HepI and HepII were performed with Clustal Omega.^38^ Resulting sequence alignment figures were made with the ENDscript server.^39^

### Molecular Modeling and Molecular Dynamic Simulation Refinement

The ternary model of HepII was made from the available apo crystal structure (PDB: 1PSW). Missing sidechains were modeled with Prime from the Schrodinger suite.^40–41^ The previously developed ternary model of HepI^42^ served as template for the placement for ADP-Hep, due to the high degree of sequence similarity occurring in the C-terminal, ADP-Hep binding domain. The C-terminal domains of HepI (residues 179-322) and HepII (residues 182-348) were aligned to aid placement of the ADP-Hep. A deacylated analog of the Hep-Kdo_2_-Lipid A acceptor substrate was docking into the center of the N-terminal cavity with a 30 Å × 30 Å × 30 Å grid box via Glide.^43^ The final docked poses were constrained to maintain contact with one of four residues (Trp10, Asp13, Lys92, Lys195) via hydrogen bonds. These residues were chosen due to their high level of sequence conservation amongst HepII homologues and their structural superposition with HepI residues that were previously determined to be in contact with its sugar acceptor. The top 5 poses according to their docking score were subjected to MMGBSA^44^ and the top two poses were selected for molecular dynamics simulations. pK_a_s for ionizable sidechains were calculated using PROPKA with 11 frames equally spaced from the second half of the trajectory. ^45–46^ Simulations were performed with the GROMACS-2021.1 simulation package and the Amber99SB forcefield.^47–49^ Ligands were prepared as previously described.^42^ Briefly, Atom types were assigned from the second generation Generalized Amber Forcefield (GAFF2) and charges were assigned with the AM1-BCC model.^50–51^ PROPKA was utilized in assigning ionization states of titratable sidechains of HepII.^45–46^ The system was placed into a dodecahedron periodic boundary condition with a 10 Å buffer region and solvated with TIP3P^52^ water model. Furthermore, the system was neutralized with the addition of 0.150 M sodium and chloride counterions. Energy minimization was performed for 50000 steps with the steepest descent algorithm. The system was equilibrated with subsequent isothermal-isochoric (NVT) and isothermal-isobaric (NPT) ensembles for a total of 12 ns with a 2 fs timestep. During equilibration, restraints (1000 kJ/mol/nm^2^) were applied to all atoms and progressively removed from the sidechains and then the backbone. Ligands were restrained during equilibration to allow the rearrangement of the active site to accommodate the ligands prior to the production simulations. Temperature and pressure were maintained with the v-rescale thermostat and Berendsen barostat with coupling of 0.1 and 2.0 ps, respectively.^48,53^ Production simulations were performed for a total of 500 ns at 300 K and 1 atm (NPT ensemble) and the temperature maintained with the v-rescale thermostat and the pressure maintained with the Parrinello-Rahman barostat with isotropic coupling.^54^ Long range electrostatics were calculated with the particle-mesh-ewald (PME) with a fourth order cubic interpolation and 1.6 Å grid spacing.^55^ Short range nonbonded interactions were calculated with a 10.0 Å cutoff. Bonds to hydrogen atoms were constrained with the LINCS algorithm.^56^ Ligand interaction diagrams were made with Maestro, molecular models were made in PyMOL and reactions were drawn in ChemDraw.^41,57^

### Expression and Purification of HepII

A pET28a plasmid containing the *E. coli* HepII (rfaF) was graciously provided by the New England Structural Genomics Consortium. The plasmid was transformed into *E. coli* BL21-AI cells (Invitrogen) and plated on Luria-Bertani (LB) agar with kanamycin (50 μg mL^−1^), with incubation for 18 hours at 37°C. One isolated colony was transferred from the plate to 10 mL of LB kanamycin (Kan) solution and incubated overnight at 37°C with shaking at 220 rpm. One liter of LB kanamycin was inoculated with 10 mL of the overnight seed culture and grown at 37°C with shaking at 220 rpm until an OD_600_ of 0.55 was reached. Protein expression was induced with a combination of isopropyl ß-D-1-thiogalactopyranoside (IPTG) and arabinose at final concentrations of 1 mM and 0.0002%, respectively. Upon induction, the temperature was reduced to 30°C and the cells were allowed to continue growing for an additional 24 hours. Cells were collected from the media by centrifugation at 5,400 × g for 10 minutes and resuspended in 20 mL of a binding buffer (20 mM HEPES, 1 mM imidazole, 0.5 M NaCl, pH 7.5) containing Lysozyme (1.25 mg mL^−1^), pepstatin A (1 μg mL^−1^) and aprotinin (1 μg mL^−1^). Cells were lysed with an Emulsiflex –C5 homogenizer at 13,000 psi and the lysate was subsequently clarified by centrifugation at 21,000 × g for 30 minutes.

A 10 mL column of Toyopearl AF Chelate 650M resin was charged with 3 column volumes (CV) of 10 mM cobalt sulfate and equilibrated with 3 CV of binding buffer (see above). Lysate was loaded at 1.0 mL/min and weakly bound proteins were eluted with an additional 3 CV of bind buffer. HepII was eluted with an imidazole gradient from a low imidazole buffer (20 mM HEPES (4-(2-hydroxyethyl)-1-piperazineethanesulfonic acid), 40 mM imidazole, 0.5 M NaCl, pH 7.5) to a high imidazole buffer (20 mM HEPES, 1M imidazole, 0.5 M NaCl, pH 7.5). Fractions containing HepII were pooled, concentrated via centrifugal ultrafiltration (30 KDa MWCO

Vivaspin 20) and buffer exchanged via Biorad P6 Desalting cartridge into a storage buffer (10 mM tris-HCl, 150 mM NaSO_4_,150 mM arginine (Arg), 40% glycerol, pH 7.5). The protein was further concentrated by centrifugal ultrafiltration (30 KDa MWCO Vivaspin 20) to 10 mg/mL, flash-frozen with liquid nitrogen and stored at −80°C. HepII can be stored in this condition for 6 months without significant loss in activity.

### Sugar Acceptor Isolation and Deacylation

*E. coli* deletion strains of HepII (ΔrfaF::Kan) and the subsequent kinase (ΔrfaP::Kan) were purchased from the Yale Keio collection.^58^ Sugar acceptor substrates were isolated from their respective knockout cell lines and prepared, as previously described.^59−60^ Briefly, two liters of cell were grown to an OD_600_ of 1 and pelleted at 5,400 × g for 10 minutes. Cells were washed with 40 mL of water, 40mL of ethanol, twice with 40 mL of acetone and once with diethyl ether. After each solvent wash, cells were pelleted at 5,400 × g and the supernatant was discarded. Cells were air dried at room temperature for one hour and pulverized with mortar and pestle prior to lipid extraction. One gram of dried cells was combined with 20 mL of a 2:5:8 phenol/chloroform/petroleum ether solution for 30 minutes while rocking. Cellular debris was pelleted at 5,400 × g for 10 minutes and supernatant was poured into a 100 mL round bottom flask. Extraction was repeated once more with the cellular debris and the resulting supernatant was post centrifugation was combined with the previously extracted supernatant in the 100 mL round bottom flask. Chloroform and petroleum ether were removed under reduced pressure. 75 mL of acetone, 15 mL of diethyl ether and 3 drops of water were added to the resulting solution to precipitate the Hep-Kdo_2_ Lipid A. The precipitant was pelleted at 20,000 × g for 10 minutes and washed with 2 mL of 80% phenol and 2 mL of diethyl ether, with the precipitate being pelleted at 20,000 × g for 10 minutes between each wash, with the supernatant being discarded. The washed precipitate was dissolved in 20 mL of water containing 0.5% triethylamine, frozen, and then lyophilized to dryness.

Hep-Kdo_2_ Lipid A was O-deacylated with hydrazine, with 10 mg of lipid reacting with 1 mL of hydrazine with stirring at 37°C for one hour. The reaction was quenched and the lipid was precipitated with 10 mL of ice cold acetone. The precipitate was pelleted at 20,000 × g for 30 minutes and the supernatant was discarded. The precipitate was washed separately with cold acetone and diethyl ether with centrifugation at 20,000 × g for 30 minutes between each wash step, with the supernatant being discarded. The washed precipitate was dissolved in 20 mL of water, frozen, and lyophilized. The formation of the O-deacylated product was confirmed via ESI mass spectrometry.

### Sugar Donor Isolation/Biosynthesis

The native sugar donor (heptosylated adenosine diphosphate, ADP-Hep) was extracted and purified from *E. coli* WBB06 cells as previously described.^29,60^ Briefly, two liters of WBB06 cells were grown in LB tetracycline (10 μg mL^−1^) to an OD_600_ of 1. Cells were collected by centrifugation at 5,400 × g for 10 minutes and the supernatant was discarded. Pelleted cells were resuspended in 40 mL of 50% ethanol (1:1 ethanol:H_2_O) for 30 minutes on ice, with stirring. The cellular suspension was centrifuged at 3,000 × g for 10 minutes to pellet cellular debris and the supernatant was kept for further processing while the pellet was discarded. Ethanol was removed under reduced pressure with a vacuum centrifuge until the volume of the sample was reduced to 20 mL. The sample was flown through an Amicon 3000 MWCO centrifugal filter via centrifugation at 3,000 × g to remove proteins that would interfere with downstream isolation and characterization of the sugar donor (ADP-Hep). The flowthrough was loaded onto a 50 mL DEAE (diethylethanolamine) column and ADP-Hep was eluted with a 0-1 M triethylammonium bicarbonate buffer (pH 8.0) gradient. Fractions containing ADP-Hep were identified via ESI (electrospray ionization) mass spectrometry. Fractions of ADP-Hep were pooled, lyophilized, and stored at −80°C. NMR data is consistent with previously published data.

The ADP-Hep analogue sugar donor ADP-mannose (ADP-Man) was biosynthesized with slight modifications as previously described.^61^ Briefly, 30 mg of mannose was combined with, 184 mg of ATP and 9.0 mg of HldE in 6 mL of a reaction buffer consisting of 500 mM tris-HCl (pH 8.0) and 5 mM MgCl_2_. The reaction was allowed to proceed overnight with monitoring via ^31^P-NMR spectroscopy. 10 μU of calf intestinal alkaline phosphatase (CIP) was added to dephosphorylate remaining ATP/ADP/AMP. The reaction mixture was then flowed over a Bio-Rad P6 desalting cartridge and lyophilized. Isolation of product was confirmed via NMR spectroscopy (^3^H, ^13^C, ^31^P).

### Circular Dichroism Spectroscopic Analysis

Protein secondary structure and thermostability were determined via Circular Dichroism as previously described for HepI.^23^ Spectra were recorded on a Jasco J-810 Spectropolarimeter and scans were obtained at varying temperatures (10°C - 95°C) between 190 nm to 250 nm in quartz cuvettes (0.2 × 0.1 cm). All conditions were performed in triplicate with 5 μM HepII in a buffer consisting of 10 mM tris-HCl (pH 7.5) and 100 mM KCl. In conditions where substrates/products (donor/acceptor) were present, 100 μM of these compounds were used to ensure complete binding of substrate to enzyme. Resulting spectra were analyzed via Matlab R2018a and melting temperature (T_M_) was calculated by fitting data to a sigmoid curve and determining the temperature at which the protein was 50% unfolded.

### Tryptophan Fluorescence Binding Studies

Emission spectra were collected on a Horiba Fluoromax-4 spectrofluoremeter. Samples were excited at 295 nm with a 6 nm slit width and resulting emission spectra was collected from 350 nm to 410 nm with a 8 nm slit width. Excitation and emission polarizers were set to 0° and 57.3°, respectively. HepII was maintained at 1 μM concentration and ligands were varied in concentration from 1 nM to 10 μM. Independent samples were made for each ligand concentration to avoid photobleaching and each concentration was repeated in triplicate. Resulting emission spectra were fit to a log normal distribution to determine the intensity at λ_max_. The intensities were then fit to a binding curve to determine the binding affinity (K_D_) with the equation 1, where [E·S] is presumed to be the fluorescence intensity that is a function of the total enzyme concentration ([E]_total_) and substrate concentration ([S]_total_)^62–63^ Resulting errors are reported as standard errors.

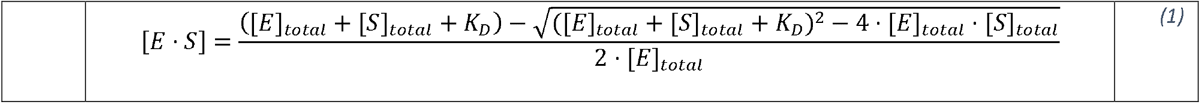

### HepII Enzyme Kinetics

Enzymatic rates were determined via pyruvate kinase/lactate dehydrogenase (PK/LDH) coupled assay as previously described.^23–24^ Each reaction mixture contained 10 nM HepII, 50 U of PK, 50 U of LDH, 100 uM phosphoenol pyruvate, 100 μM NADH and 100 μM dithioerythritol in a reaction buffer (50 mM HEPES, 50 mM KCl, 10 mM MgCl_2_, pH 7.5). When varying one substrate, the other substrate was introduced at 10,000 fold excess (100 uM). The sugar donor/acceptor were varied between 100 nM to 20 μM. Reduction of NADH was monitored at 340 nm with a constant temperature of 37° C on a Cary UV-Vis spectrophotometer. Reactions were initiated by the addition of enzyme. Reactions involving the lowest concentration of substrate (100 nM) approached the sensitivity limit of the instrument, therefore, initial rates were determined by fitting the full progress curve rather than the first 10% with equation 1.^64^ The Michaelis constant (K_M_) and turnover number (*k_cat_*) were calculated by fitting initial rates as a function of substrate concentration to the Michaelis-Menten equation.

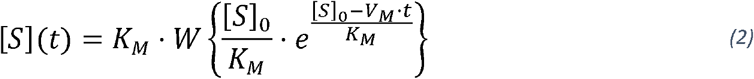

## Results

### Multiple Sequence Alignment and Sequence Conservation Analysis

HepII catalyzes the transfer of a heptose onto the growing LPS inner core region (Figures 1-2). This enzymatic reaction is directly preceded by HepI that similarly transfers a heptose residue onto the Kdo_2_-Lipid A, producing the substrate for HepII (Figure 1). Based upon previous molecular biology experiments, both heptosyltransferase enzymes use an ADP-L-*glycero*-D-*manno*-heptose (ADP-Hep) as their sugar donor substrate. A sequence similarity comparison between *E. coli* HepI and HepII reveals a 32% sequence identity, with common residues located mainly within the C-terminal Rossmann-domain, where the ADP-Hep is expected to bind (Figure 3, S1). Furthermore, as previously observed in a multiple sequence alignment of HepI-III, they share a highly conserved Asp13 that is believed to facilitate the transfer reaction by acting as a base during the sugar transfer reaction (Figure 2, S1-S2).^25,65^ To elucidate the residues potentially involved in HepII substrate interactions, we performed a sequence alignment of HepI with HepII (HepI PDB: 6DFE and HepII PDB:1PSW; Figure S1), in addition to generating representative multiple sequence alignments (MSAs) of HepI (Figure S2) and HepII (Figure 4) homologues. Previously identified residues in HepI that are involved in substrate binding are indicated^21,25,42^ and were used as a guide to identification of residues in a HepII that might be important for catalysis, in addition to examining residues that are highly conserved in a multiple sequence alignment (MSA) of HepII. Based upon these analyses of HepII, we hypothesized FDHLA binding residues to be Pro8, Trp10, His61, Ser90, Lys92, Gly192, Lys195, and Asp269 (Table 1A, S1). For the native sugar donor, HepII potentially important residues involved in ADP-Hep binding include Trp10, Glu190, Gly192, Lys195, Thr250, Leu252, Ala255, Asp269, Ser270, Gly271, Leu272, and His274 (Table 1B, S1). In our multiple sequence alignment of HepII (Figure 4), a high degree of conservation of Asp13 in HepII homologues from a variety of species was observed, which is consistent with observations that Asp is a conserved catalytic base for inverting GT-B enzymes.^66^ Furthermore, each of the HepII residues listed above maintain at least 90% conservation in a representative MSA generated from 150 HepII homologues spanning 35-95% identity to the HepII from *E. coli*.

**Table 1.**
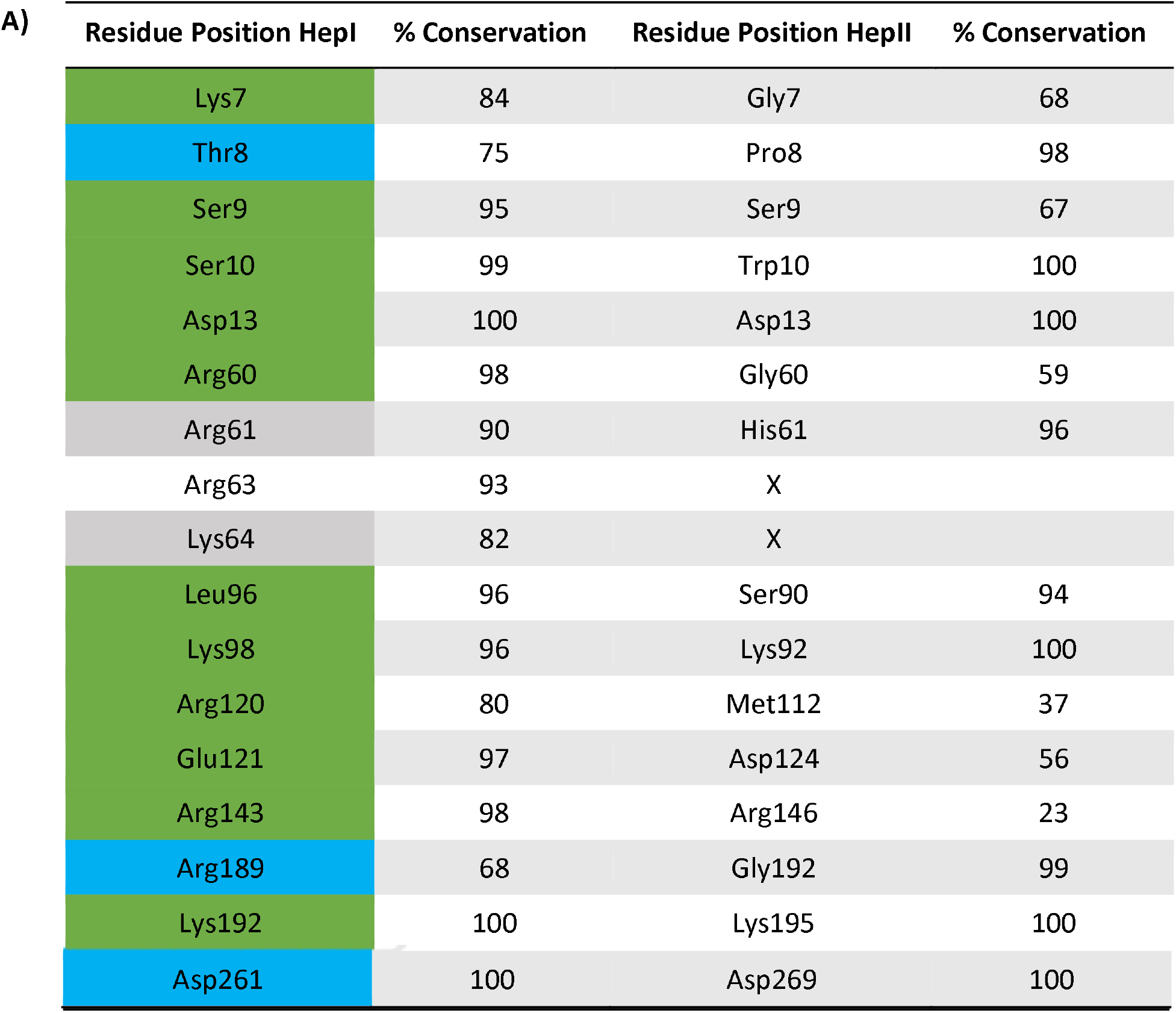
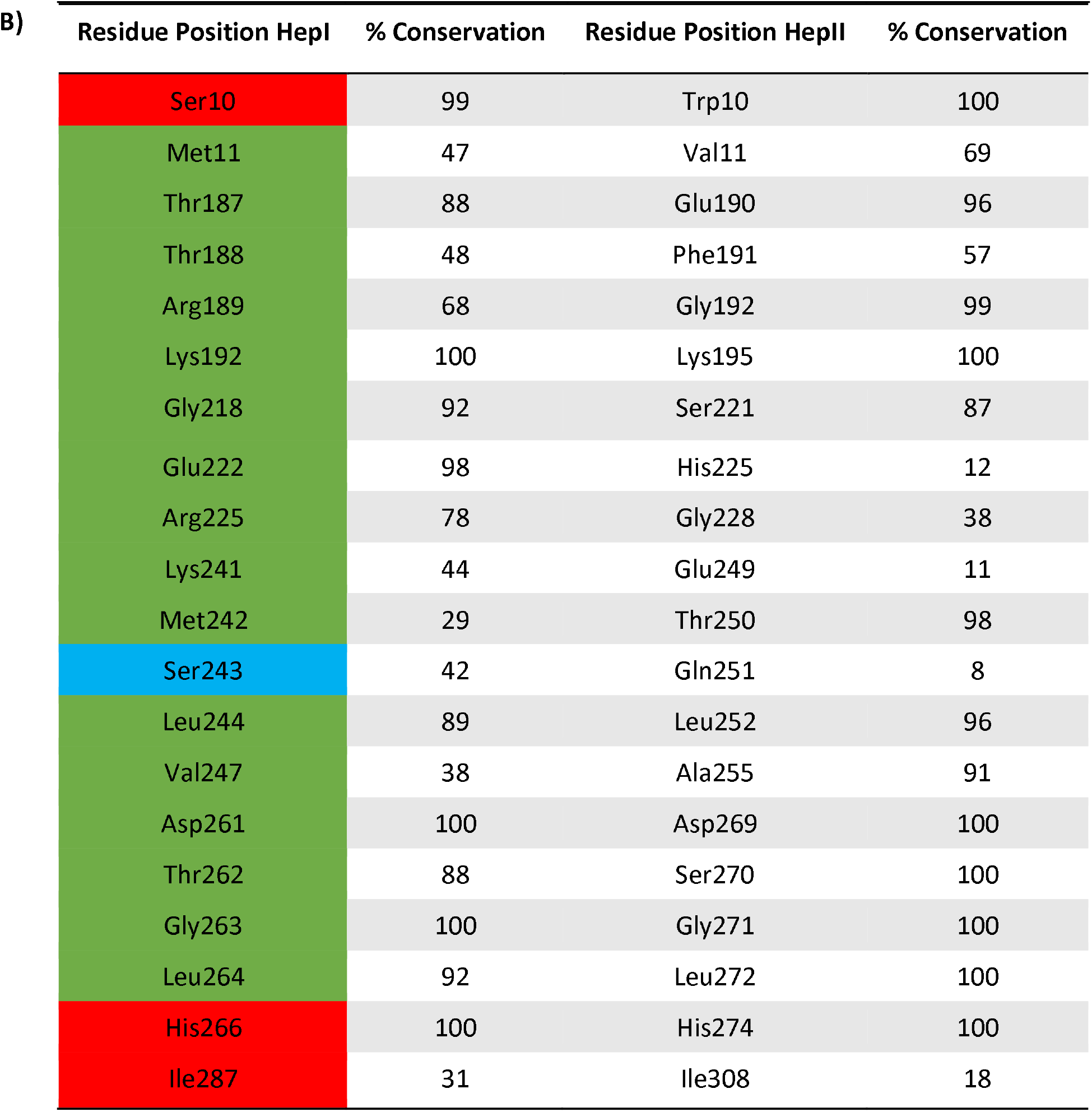
Conserved binding residues in HepI and equivalent residues in HepII. HepII residues equivalent to HepI binding residues based on multiple sequence alignment for **(A)** FDLA/FDHLA and **(B)** ADP-Hep/ADP. HepI residues are color coded by contact with substrate (red), product (cyan), or both (green).

**Figure 4.**
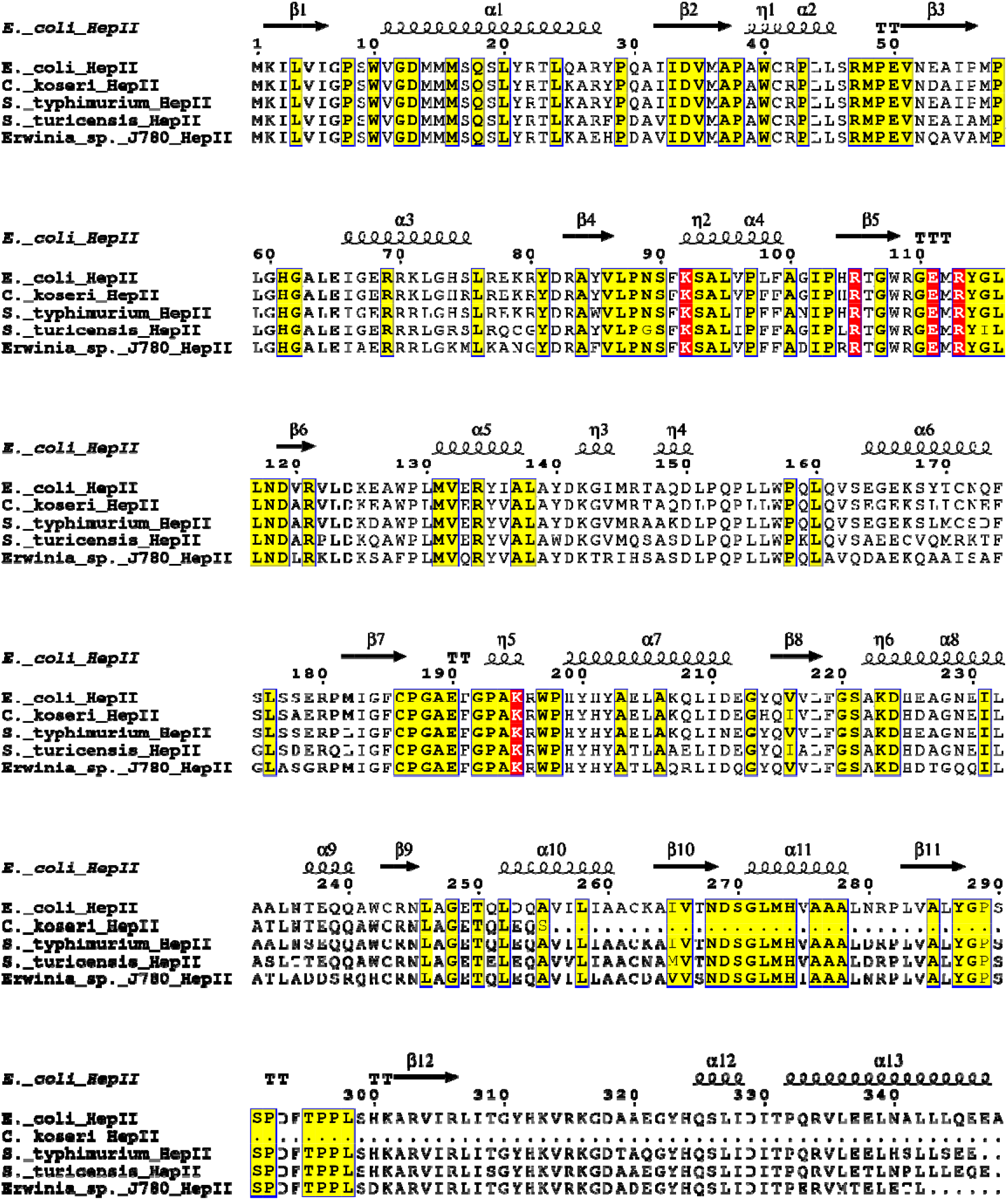
Representative sequences of HepII multiple sequence alignment. First 5 sequences of multiple sequence alignment for HepII with the query sequence (first sequence) from the crystalized E. coli HepII (PDB: 1PSW) and the secondary structural elements of the first sequence are represented immediately above the sequence. Residues in red boxes denote strictly conserved residues, and ones in bolded/boxed are greater than 70 similar.

### Ternary Model Development and Refinement

Both HepI and HepII are part of the GT-B structural class, and a structural alignment yields a superposition with a C_α_ root mean square deviation of their structures of 3.8. Due to the high sequence similarity of the C terminal domains of HepI and HepII, placement of the sugar donor (ADP-Hep) in HepII was modeled (Figure 3) through a superposition of previously observed crystal structure orientation of the complex ADP-Hep with HepI (PDB 2H1H). For the sugar acceptor substrate (FDHLA), we docked this ligand in the N-terminal domain of HepII, so as to maintain a minimum of one hydrogen bond contact with each highly conserved residues identified from the MSA analysis predicted to be involved with FDHLA binding, including residues Trp10, Asp13, Lys92, Lys195 (Table 1A). In pose2, there is electrostatic complementarity in the active site with arginines and lysines in the active site (Figure 3). The top two poses were subjected to further structural minimization via 500 ns molecular dynamics (MD) simulation. The apo form was also simulated to provide a reference for the unbound complex.

The backbone root mean square deviation (RMSD) across the second half of the trajectory for apo, pose1, pose2 complexes are 2.08 ± 0.16 Å, 2.12 ± 0.18 Å, 1.88 ± 0.10 Å, respectively (Figure 5A, S11A, Table S3). The C^α^ root mean square fluctuations (C”RMSF) for apo and pose1 reveals residues with fluctuations greater than 1.5 Å in the 60s and a majority of the C-terminal domain (Figure S11B, Table S3). Whereas pose2 only has a 15 residues with fluctuations greater than 1.5 Å in the 230s and 300s (Figure 5B, S11B, Table S3). To assess the merits of the predicted ligand binding residues from the MSA examination (above), the HepII•ADP-Hep•FDHLA ternary complex simulation was used to identify amino acids adjacent to the ligands and ligand interaction maps were generated (Figure 6). In the simulation of the ternary complex (pose2), the acceptor (FDHLA) is held in place by electrostatic interactions between Arg69, Lys92, Lys316 and the phosphate groups on the acceptor (Figure 6A, S12A). The acceptor is further stabilized by hydrogen bonding interactions with other residues including Ser9, Trp10, and Asn89. Interactions between the acceptor and HepII in pose1 only differs by an electrostatic interaction, where the Lys316 of pose2 is replaced by Lys125 in pose1 (Figure S13A, S14A). The donor (ADP-Hep) is stabilized by electrostatic contact between the alpha phosphate and Lys195 (Figure 6B, S12B). Furthermore, a hydrogen bonding interaction between the primary amine of the adenosine ring and Thr250 sidechain aids in holding the donor in place. There is also a T shaped pi-pi stacking interaction between Trp40 and the indole ring of the adenine. Lastly, there are additional hydrogen bonding interactions between ADP-Hep and residues Glu190, Gly271, Asp269, His 274 and Lys313. In pose1, very similar interactions are observed and only differing by the lack of pi-pi or hydrophobic interactions between the acceptor and Trp40 (Figure S13B, S14B).

**Figure 5.**
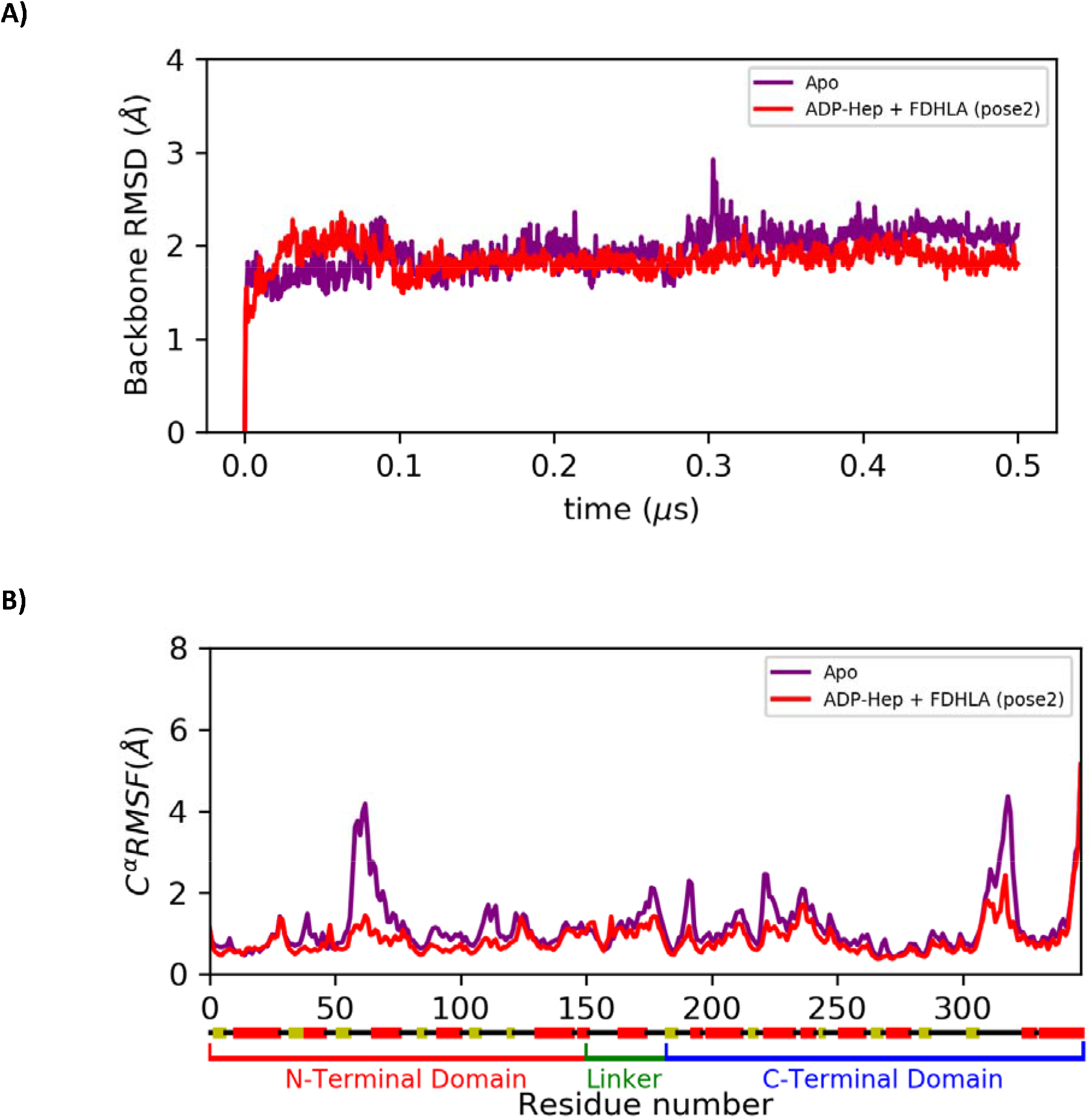
RMSD and RMSF of HepII trajectory. (A) Backbone RMSD and (B) Cα RMSF of HepII apo and pose2 of the HepII•ADP-Hep•FDHLA complex.

**Figure 6.**
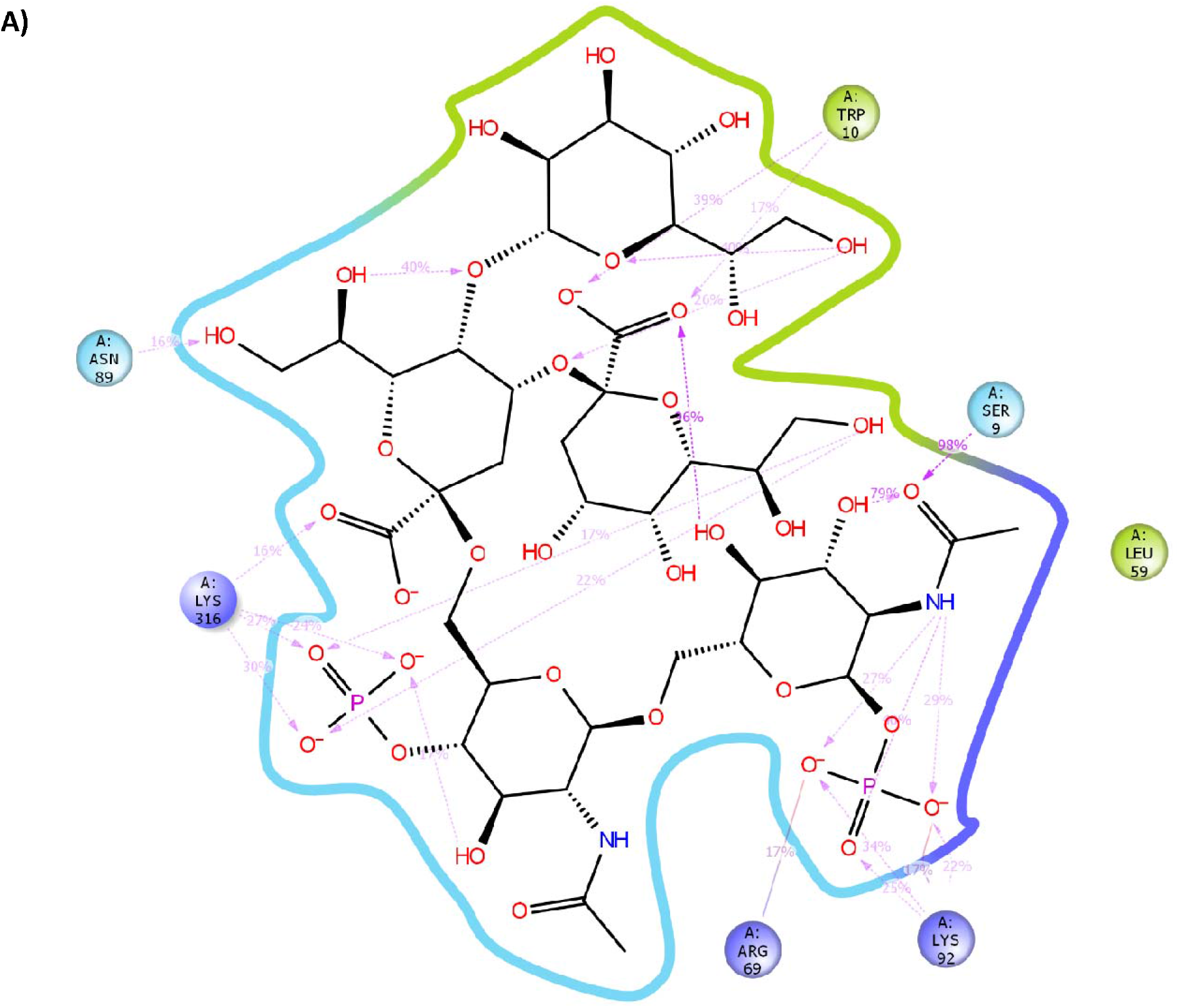
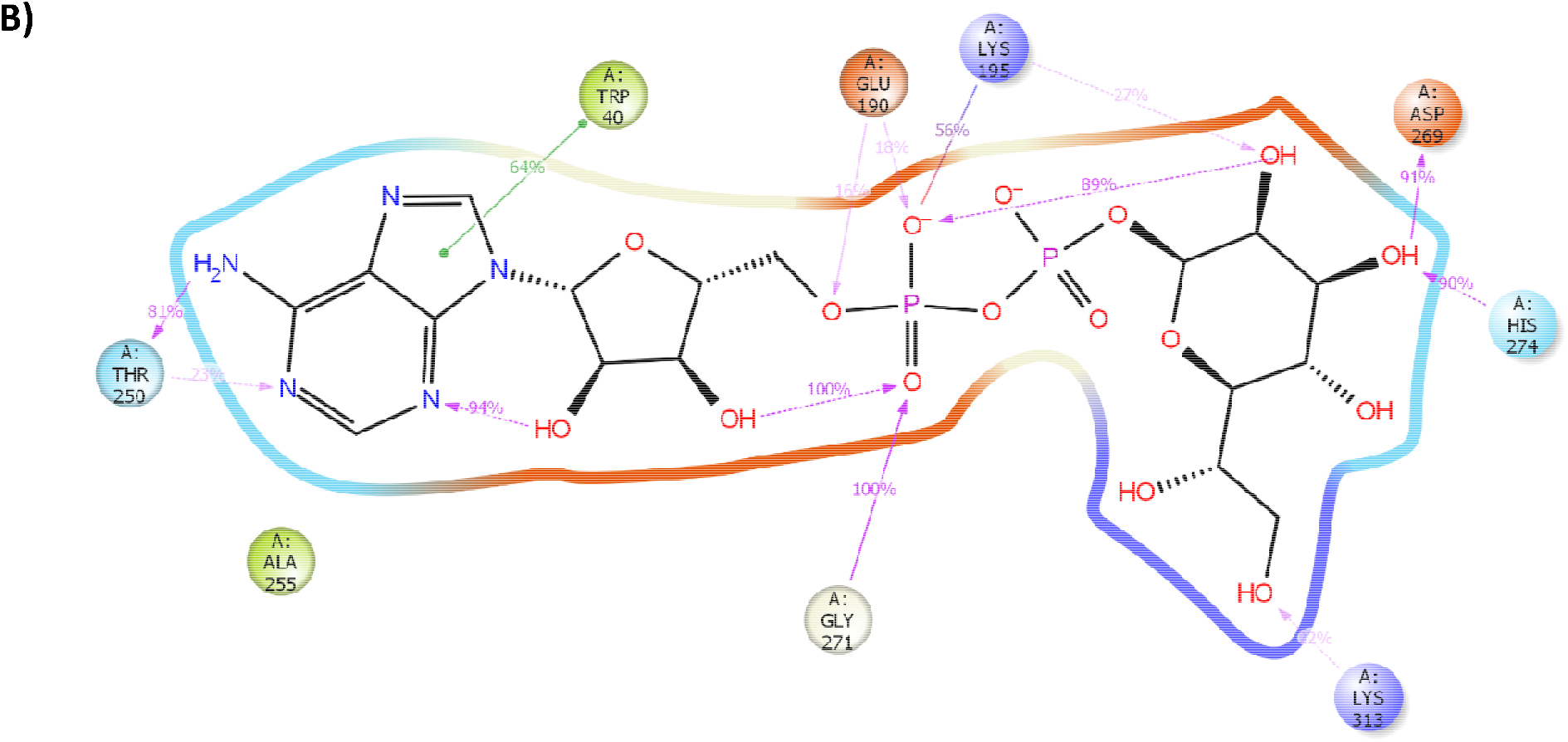
Ligand interaction diagram of substrates to HepII sidechains across MD trajectory. (A) Contacts between acceptor substrate, FDHLA (pose2), and lysine/arginine of HepII that anchor ligand in N-terminal side of active site interface. (B) Lysine and tryptophan sidechains that anchor donor, ADP-Hep, in C-terminal side of active site interface as observed from MD trajectory.

Lastly, the pK_a_ of ionizable sidechains were calculated to probe whether the catalytic residue has a perturbed pK_a_ in-line with the hypothesized mechanism of the transfer reaction (Figure 2). The pK_a_ of Asp13 in HepII apo, ternary complex pose1 and pose2 are 5.31 ± 0.46, 5.26 ± 0.71, and 6.17 ± 0.32 (Table S4). A similar elevation of the pK_a_ of Asp13 in HepI was previously observed.^42^

### Experimental Secondary Structural Characterization of HepII

HepII from *E. coli* was successfully purified to greater than 90% purity (Figure S3). The secondary structural characteristics of HepII, as determined by CD spectral analysis, suggest a protein with a mixed α/β structure in the apo state (Figure 7A). In the presence of ODHLA, HepII gains a more pronounced second minima, suggestive of an increase in alpha helicity. The melt temperature of HepII apo is 60° C (Figure 7B, S4A). In the presence of the sugar acceptor (ODHLA), HepII maintains secondary structure at higher temperatures (95° C) with an increase in the overall signal intensity at the same protein concentration (Figure 7B, S4A-B). The sugar acceptor product (ODH_2_LA) has similar stabilizing affect and induces the same secondary structural changes as ODHLA, but with a decrease in the signal intensity at the same protein concentration (Figure 7B, S7E). ADP and ADP-Hep have similar melt temperatures and do not exhibit any substrate induced stabilizing affect or secondary structure change as with ODHLA or ODH_2_LA (Figure 7B, S7B-D).

**Figure 7.**
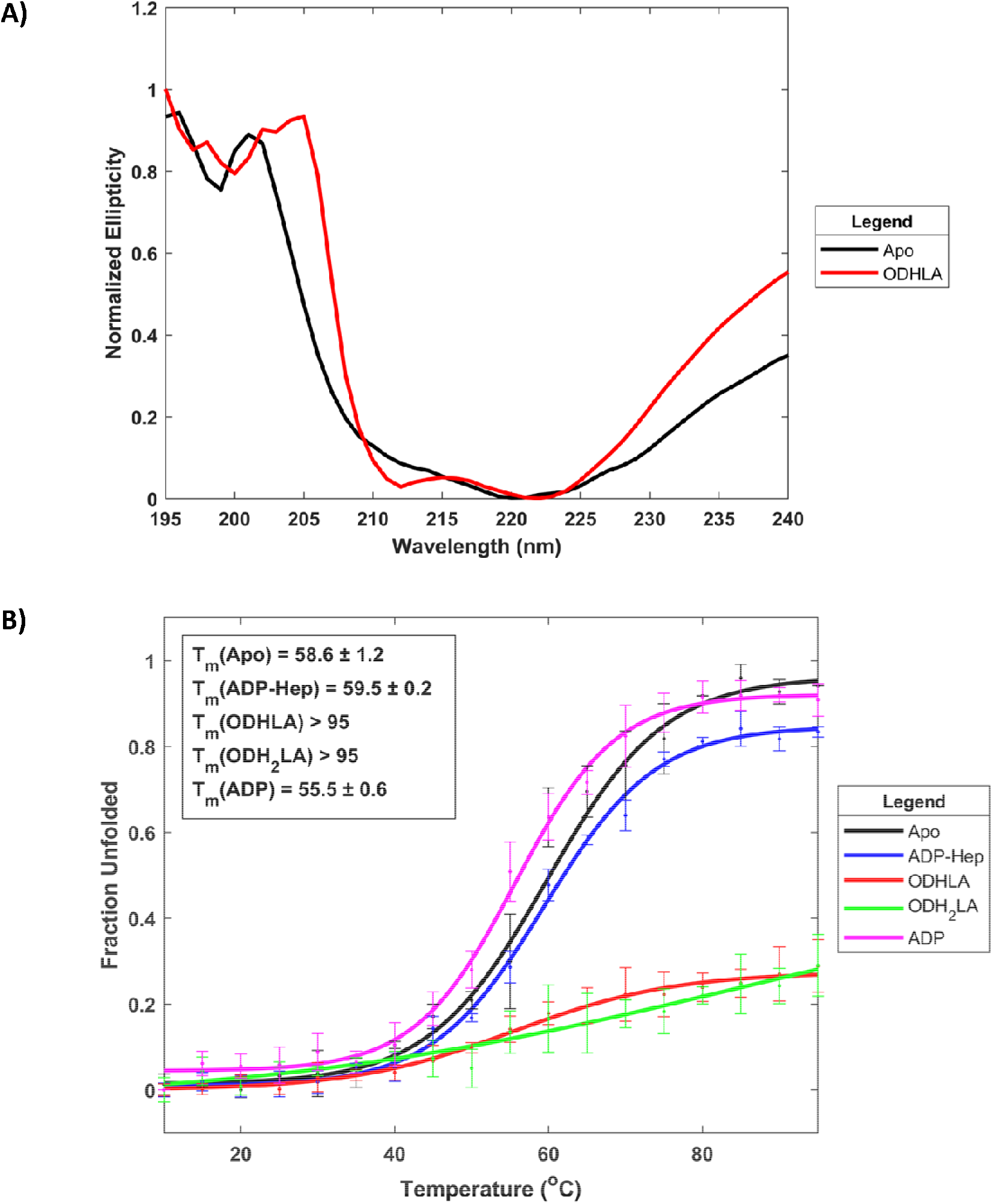
Circular dichroism spectroscopy of HepII with and without ligands and at various temperatures. **(A)** Secondary structure profile of HepII in the absence/presence (black/red) of ODHLA (acceptor) and demonstrates an increase in helicity due to a deepening of the second minima at 210 nm. **(B)** Unfolding of HepII as a function of temperature with substrates/products as determined by signal intensity at 220 nm demonstrates stabilization of HepII at high temperatures in the presence of the acceptor substrate/product (red/green) relative to apo (black).

### HepII Substrate Binding and Kinetics

HepII has seven tryptophan residues that can act as potential reporters for binding. As mentioned above, Trp10 is in the active site and sits in the space between both ligands and is highly conserved (Table 1). In the presence of the native sugar donor (ADP-Hep) or acceptor (ODHLA), HepII’s tryptophan fluorescence emission maximum exhibits a decrease in signal intensity as a function of concentration as compared to that of the apo protein (Figure 8A, S5). Using this change in fluorescence intensity upon ligand binding, binding affinities of native substrates to HepII were determined. The native sugar donor (ADP-Hep) has a *K*_D_ of 0.66 ± 0.46 μM (Figure 8B, S5, Table 2). For the sugar acceptor substrate analog (O-deacylated-Hep-Kdo_2_ Lipid A) has a *K*_D_ of 0.48 ± 0.14 μM (Table 2, Figure S5).

**Figure 8.**
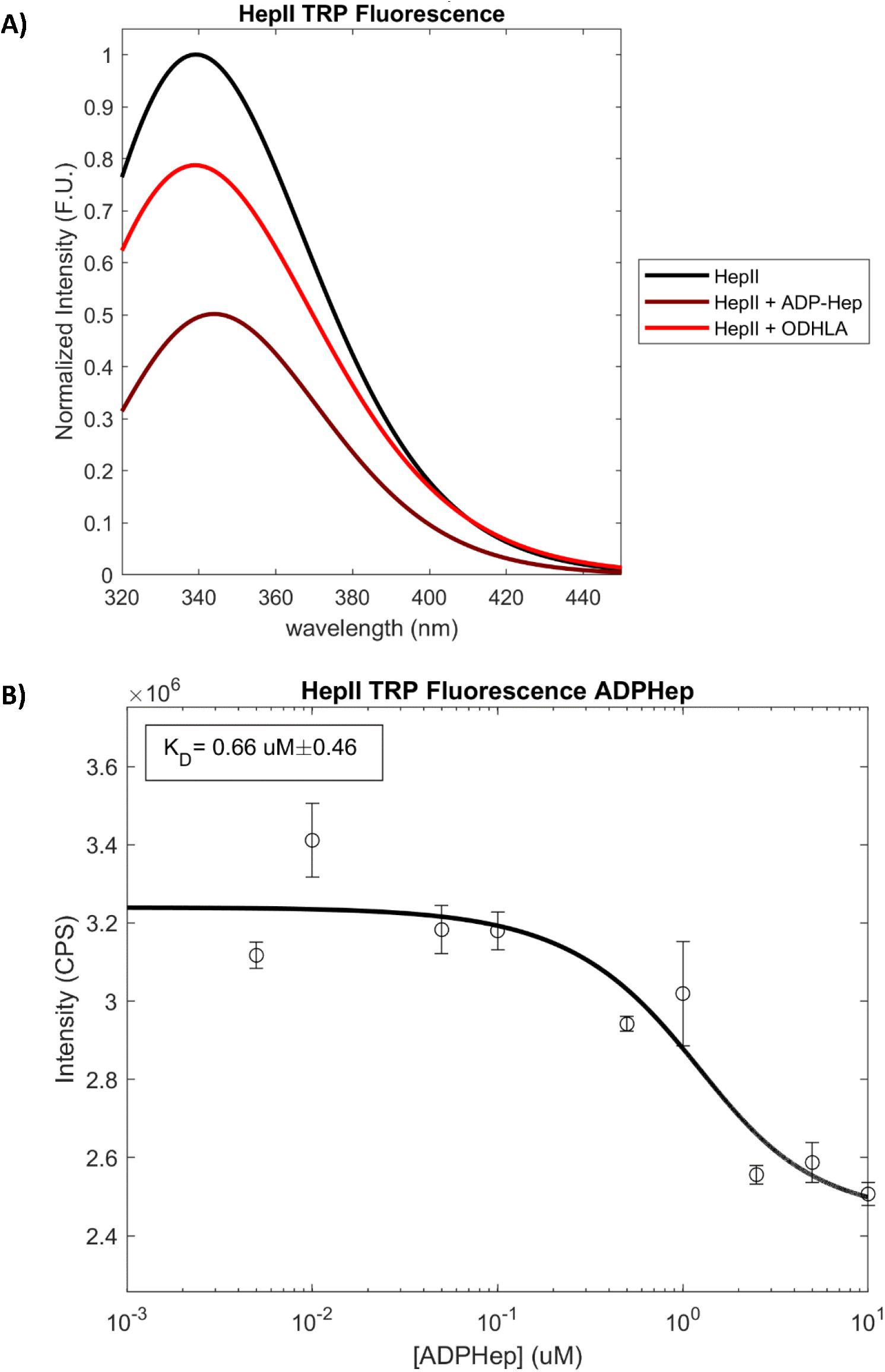
Tryptophan fluorescence emission spectra and ligand binding curve. **(A)** Overlay of HepII Tyrptophan fluorescence emission spectra of HepI apo (black), with ADP-Hep (brown), and ODHLA (red). **(B)** Binding curve of ADP-Hep as determined by quenching of tryptophan fluorescence as a function of APD-Hep concentration.

**Table 2.** Michaelis constant, turnover number, catalytic efficiency, and binding affinities of HepII substrates. Kinetic parameters determined for HepII for fully acylated (H-Kdo2-Lipid A) and partially deacylated (O-deacylated-H-Kdo2-Lipid A) donor and acceptors (ADP-Hep, ADP-Man) and binding affinities as determined via tryptophan fluorescence quenching.

| | $k_{cat} (s^{-1})$ | $K_M (\mu M)$ | $k_{cat}/K_M (s^{-1} M^{-1}) \times 10^6$ | $K_D (\mu M)$ |
| --- | --- | --- | --- | --- |
| **HLA w/ Triton X 100** | 0.57 ± 0.06 | 0.49 ± 0.13 | 1.2 ± 0.4 |  |
| **HLA** | 0.33 ± 0.03 | 0.41 ± 0.09 | 0.8 ± 0.2 |  |
| **ODHLA** | 0.423 ± 0.007 | 0.58 ± 0.08 | 0.7 ± 0.1 | 0.48 ± 0.14 |
| **ADP-Hep** | 0.47 ± 0.02 | 0.57 ± 0.09 | 0.8 ± 0.2 | 0.66 ± 0.46 |
| **ADP-Man** | 0.54 ± 0.02 | 0.77 ± 0.08 | 0.7 ± 0.1 |  |

To gain further insight into HepII catalysis, the turnover rate and Michaelis equilibrium constant were determined for the native/non-native sugar donors and acceptors. For the native sugar acceptor, the kinetics were determined in the presence and absence of Triton X-100 to ensure micelle formation of the acceptor was not inhibiting the reaction (Table 2, S1, Figure S6). For HLA with (without) triton X-100, the *k_cat_* and *K*_M_ are 0.57 ± 0.06 s^−1^ (0.33 ± 0.03 s^−1^) and 0.49 ± 0.13 μM (0.41 ± 0.09 μM), respectively. The O-deacylated-Heptosyl-Kdo_2_ Lipid A has a *k_cat_* of 0.423 ± 0.007 s^−1^ and *K*_M_ of 0.58 ± 0.08 μM. When varying sugar donor concentrations, the ADP-Hep (ADP-Man) *k_cat_* and *K*_M_ are 0.47 ± 0.02 s^−1^ (0.54 ± 0.02 s^−1^) and 0.57 ± 0.09 μM (0.77 ± 0.08 μM), respectively.

## Discussion

### Conserved Binding Site Residues

HepI and HepII facilitate consecutive transferase reactions with an identical sugar donor (ADP-Hep) and a sugar acceptor where the product of the HepI reaction is the substrate of the HepII reaction (Figure 1-2). It is therefore not surprising that most of the sequence conservation between these two enzymes occurs in the C-terminal domain (Figure S1). More surprising is the lack of sequence conservation in the N-terminal domain to facilitate binding of HLA which is a common product/substrate between the two enzymes. Examination of conserved active site residues reveals maintenance of the catalytic base and three of nine positively charged arginine (Arg) and lysineslysine (Lys) residues that in HepI are responsible for anchoring the negatively charged phosphate groups on the acceptor via electrostatic interactions. Only the homologues of HepI residues Lys98, Arg 143 and Lys192, are maintained and conserved HepII (corresponding to residues Lys92, Arg146 and Lys195 in HepI). Arg146 is greater than 20 Å from the nearest phosphate on the ligand, therefore, it may not play a major role in substrate binding and stabilization. The Lys195 stabilizes the donor and is unavailable to bind to the acceptor in the C-terminal domain (Figure 6). Charged residues that bind to the acceptor, suggested by our simulations include Arg69, Lys125, Lys313, Arg315, and Lys316. Residues Lys313, Arg315 and Lys316 are conserved 17%, 24% and 57% of the time, respectively. Therefore, they are likely to play a minor role in substrate binding. Arg69 is conserved 98% of the time, therefore, it may coordinate to the same phosphate as Lys92 (Figure 6A). Lys125 is conserved 73% of the time and is a better candidate for anchoring the acceptor substrate in the active site. We hypothesize that these conserved charged residues may facilitate binding of the acceptor substrate and subsequent promotion of conformational changes that are necessary for catalysis by bringing the nucleophilic hydroxyl of the acceptor close to the anomeric carbon of the donor (Figure 6A, S12A, S14A), as has been previously observed in HepI.^42^ Both enzymes have a highly conserved Asp13 that potentially acts as the catalytic base that removes the proton from the hydroxyl group of the acceptor to promote a nucleophilic attack on the anomeric carbon (Figure 2). This is further suggested by the perturbed pK_a_s of Asp13 calculated for HepII and previously calculated for HepI (Table S4).^42^ HepI has a serine at the 10^th^ position, whereas, HepII has a tryptophan (Trp), so no other potentially conserved catalytic residues have been identified. Due to the added sugar on the HepII substrate, the stabilizing hydrogen bond from the serine in HepI may have been replaced by a tryptophan to form hydrophobic interactions with the hydrophobic face of this added sugar. Since Trp10 also is predicted to be adjacent to the bound ADP-Hep, this residue could be important for form anion-π interactions^67^ with the phosphates of ADP-Hep. Alternatively, as suggested by our modeling, the highly conserved Trp40 (92%) may be interacting with the indole ring of the donor and contributes to the fluorescence quenching that is observed experimentally for the donor, while the substrate induced quenching of the acceptor is due to hydrophobic interactions with the acceptor (Figure 6). In addition to the Trp10, HepII has several other HepI equivalent residues that similarly stabilize ADP-Hep binding. The backbone of Glu190 amine forms hydrogen bonds with the phosphates of ADP-Hep, similar to the HepI equivalent Thr187. The HepI residue Met 242 which is strictly conserved and responsible for binding of the NH_2_ group of the adenine base through a backbone carbonyl, is now replaced by Thr250, which is conserved 98% of HepII homologues analyzed. Furthermore, HepII seems to have a highly conserved heptose “sensing” residue Asp269 that hydrogen bonds to the hydroxyl groups of the heptose moiety, analogous to Asp261 in HepI. Overall, examination of the sequence and structural superpositions of HepI and HepII have allowed identification of a collection of amino acid residues that warrant further investigation for their involvement in ligand binding and catalysis in subsequent studies.

### Binding Induced Structural Changes

In the crystal structure of HepII, there are two stretch of residues that are disordered (58-64, 311-320). The flexibility of these two regions are evident in the RMSF including downstream residues 65-77 (Figure 5B, S11B). Residues 58-64 are analogous to residues 58-70 in HepI which have been previously observed to adopt a more alpha-helical orientation upon ligand binding and has been observed via CD and through crystal structures (Apo: PDB 2GT1, Psuedoternary: PDB 6DFE). ^21, 23, 25, 42^ In our HepII simulations, residues 58-64 become less dynamic upon presence of the acceptor in both poses (Figure 5B, S11B) which could promote formation of an ordered secondary structure. Additionally, the disordered 60s loop adds a turn to the downstream helix and the helix gains a turn on either side (Figure S15). This *in silico* observation of a ligand induced secondary structural rearrangement is also supported by observed changes in the CD spectral comparisons of HepII apo and liganded forms (Figure 7A). These combined observations suggest that HepII experiences a net increase in α helicity in the presence of the sugar acceptor and this could be attributed to the stabilization of the 60s α helix downstream of the disordered loop (58-64) (Figure 7A, S15). Furthermore, residues Lys195 in the C-terminal domain also becomes ordered by the presence of the sugar acceptor, whereas the acceptor binds in the N-terminal domain. Engagement of both domains by the acceptor may explain why only in the presence of the acceptor does HepII experience significant thermal stabilization.

Based on our ternary structural model of HepII, we hypothesized that binding of either substrate would have an impact on the tryptophan fluorescence emission spectra because of the proximity of Trp10 or Trp40 to both ligands (Figure 8A). Indeed, we observe a concentration dependent quenching of tryptophan fluorescence upon binding of both the sugar donor and the sugar acceptor. We hypothesize that this tryptophan may be there to assist in the exclusion of water from the active site by acting as part of a cage surrounding the Asp13, the acceptor hydroxyl and the anomeric donor. This tryptophan may also be important in intercalating into the membrane to enhance acceptor substrate binding of HepII to enable catalysis. Our modeling also suggests a possible role for Trp40 aiding in substrate binding of the donor. Further studies are needed to definitively understand the mechanistic role of Trp10 and Trp40, including studies of HepII binding to a membrane anchored acceptor substrate embedded in a micelle, to elucidate its role (if any) in catalysis.

### Substrate Selectivity Evaluation

While the enzyme that catalyzes the Kdo-transfer reaction early in the inner-core LPS biosynthesis is sensitive to the number of acyl chains on the acceptor^68^ the same is not true of HepI or HepII. The deacylation of the HepI and HepII sugar acceptors provide tremendous practical advantages for experiments by increasing the solubility of the substrate and eliminating the need for detergents. As previously demonstrated, HepI experiences a 2-5 fold increase in the catalytic efficiency upon deacylation of the acceptor mainly due to a decrease in the *K*_M_.^24^ On the contrary, the catalytic efficiency of HepII is unaffected by the presence or absence of acyl groups on the acceptor where an increase in the turnover rate (*k_cat_*) is counterbalanced by an increase in the *K*_M_ (Table 2, Table S1). Furthermore, as previously demonstrated *in vivo^26^* and quantified in our work, HepII has an approximately 2.3 fold increase in its catalytic efficiency due to a greater affinity for the acceptor substrate as compared by the differences in the *k_M_*. Additionally, HepII can also utilize a hexose (ADP-Man) sugar donor, as previously observed in HepI. The *K_M_* for the hexose is approximately 1.4 fold lower than the native heptose, again being less impacted than HepI which exhibits a nearly 10 fold decrease in the Michaelis constant for the same reduction in susbtrate size.^69^ The binding affinities determined by tryptophan fluorescence quenching yield similar affinities for O-deacylated acceptor and native donor. Overall, the substrate utilization profiles for the two enzymes are quite similar and the ability of HepII to also utilize a sugar acceptor substrate with fewer acyl chains has the potential to allow for future work to obtain liganded crystal structures of HepII to validate the predicted ligand-binding interactions described here.

## Conclusion

Our work here has provided the first quantitative determination of HepII functionality and its similarities and differences with HepI. In light of the low sequence similarity between HepI and HepII, their structural homology and multiple sequence alignments have allowed us to infer the conserved residues that may contribute to substrate binding and catalysis. This information has also allowed us to create a working model for the ternary complex HepII, as previously done for HepI.^42^ This working model inspired the use of tryptophan fluorescence to determine the binding affinities of native and non-native donor and acceptor ligands, which also helps substantiate the validity of the ternary structural model of HepII. HepII has a higher catalytic efficiency and a submicromolar affinity for its substrates, which is distinct from prior observations for HepI. Further studies will need to be performed to fully understand the importance of these differences in kinetic parameters. HepII’s insensitivity to the number of acyl chains on the acceptor may be a result of the longer sugar polymer region and suggests that HepII may not need to rely on the lipids as another source of ligand contacts. This result coupled with the smaller *K*_M_ for HepII binding of the sugar acceptor substrate could enable HepII to be a better target for inhibitor design studies, but further research is needed to determine if tight-binding inhibitors can be developed for this enzyme. Additionally, changes in secondary structure and thermal stability in the presence of the acceptor is akin to HepI and may be a conserved behavior for heptosyltransferase enzymes, or perhaps GT-B enzymes more broadly. This work presents new insights and confirms previously determined *in vitro* characteristics of HepII that paves the way for the future design of novel therapeutics against this family of enzymes.

## Accession codes

More information available about Heptosyltransferase II from *E. coli* can be found in the PDB (PDB ID: 1PSW) and Uniprot databases (ID: P37692).

## Abbreviations

(GT): glycosyltransferase
(HepII): heptosyltransferase II
(Hep): L-*glycero*-D-*manno*-heptose
(ADP-Hep): ADP-L-*glycero*-D-*manno*-heptose
(LPS): lipopolysaccharide
(HLA): *E. coli* Hep-Kdo_2_-Lipid A
(ODHLA): O-deacylated *E. coli* Hep-Kdo_2_-Lipid A
(Trp): tryptophan
(Phe): phenylalanine
(CD): circular dichroism
(MD): molecular dynamics
(TLR4): toll-like receptor 4
(CaZY): Carbohydrate-Active enZYme
(ADP-Man): ADP-β-mannose
(MMGBSA): molecular mechanics generalized Born surface area
(GAFF2): Generalized Amber Forcefield
(LB): Luria-Bertani
(CV): column volume
(PK/LDH): Pyruvate Kinase/Lactate Dehydrogenase
(MSA): multiple sequence alignment
(T_M_): melting temperature

## Acknowledgements

Daniel J. Czyzyk and Noreen K. Nkosana for assistance in determining the conditions for HepII protein expression, the New York Structural Genomics Research Consortium for a plasmid containing the subcloned *E. coli* HepII gene and the Minnesota Super Computing Institute for the Schrodinger Suite.

## Notes

### Competing Interest Statement

The authors have declared no competing interest.

