## Supplemental Information for "Kinetic Characterization and Computational Modeling of the *Escherichia coli* Heptosyltransferase II: Exploring the Role of Protein Dynamics in Catalysis for a GT-B Glycosyltransferase"

Table S1: **Michaelis constant, turnover number, and catalytic efficiency of HepII substrates.** Kinetic parameters determined for HepII for fully acylated (H-Kdo_2_-Lipid A) and partially deacylated (O-deacylated-H-Kdo_2_-Lipid A) donor and acceptors (ADP-Hep, ADP-Man). Left most column indicates the substrate that is varied and the and the top row indicates which substrate was introduced in excess.

|  | ADP-Hep | | | ADP-Man | | |
| --- | --- | --- | --- | --- | --- | --- |
|  | *k_cat_* (s^-1^) | *K_M_* (μM) | *k_cat_*/*K_M_* (s^-1^ M^-1^) x 10^6^ | *k_cat_* (s^-1^) | *K_M_* (μM) | *k_cat_*/*K_M_* (s^-1^ M^-1^) x 10^6^ |
| HLA w/ Triton X 100 | 0.57 ± 0.06 | 0.49 ± 0.13 | 1.2 ± 0.4 | 0.33 ± 0.03 | 0.52 ± 0.11 | 0.6 ± 0.2 |
| HLA | 0.33 ± 0.03 | 0.41 ± 0.09 | 0.8 ± 0.2 | 0.78 ± 0.05 | 0.50 ± 0.08 | 1.6 ± 0.4 |
| ODHLA | 0.423 ± 0.007 | 0.58 ± 0.08 | 0.7 ± 0.1 | 0.44 ± 0.07 | 0.54 ± 0.17 | 0.8 ± 0.4 |
|  | ODHLA | | | | | |
|  | *k_cat_* (s^-1^) | | *K_M_* (μM) | | *k_cat_*/*K_M_* (s^-1^ M^-1^) x 10^6^ | |
| ADP-Hep | 0.47 ± 0.02 | | 0.57 ± 0.09 | | 0.8 ± 0.2 | |
| ADP-Man | 0.54 ± 0.02 | | 0.77 ± 0.08 | | 0.7 ± 0.1 | |

Table S2: **MMGBSA and glide scores of docked poses.** Glide scores and binding energies (MMGBSA) of docked acceptor (FDHLA) to HepII•ADP-Hep complex.

| **FDHLA Docking Pose** | **Glide Score** | **ΔG_MMGBSA_ (kcal/mol)** |
| --- | --- | --- |
| Pose 1 | -6.205 | -28.07 |
| Pose 2 | -5.450 | -1.57 |
| Pose 3 | -5.395 | -23.71 |
| Pose 4 | -5.143 | -20.45 |
| Pose 5 | -5.090 | -14.48 |

Table S3: **RMSD and RMSF values of HepII simulations.** Average quantities of backbone RMSD and residues with fluctuations greater than 1.5 Å.

| **Complex** | **RMSD (Å)** | **RMSF > 1.5 Å** |
| --- | --- | --- |
| HepII | 2.08 ± 0.16 | 58-71, 111-115, 174-180, 191-193, 211-213, 222-227, 233, 236-241, 310-322 |
| HepII•ADPHep•FDHLA (pose1) | 2.12 ± 0.18 | 59-67, 114-115,126-127,149-150,152-154,172-180,191-192,211,213,226,229,233,236-242,293-294,308-322 |
| HepII•ADPHep•FDHLA (pose2) | 1.88 ± 0.10 | 236-238, 310-313, 316-318 |

Table S4: **Average pK_a_ of D13 from MD simulations.** pK_a_ of putative catalytic sidechain as determined by PROPKA.

| **Complex** | **Residue** | **pK_a_ (PROPKA)** |
| --- | --- | --- |
| HepII | Asp13 | 5.31 ± 0.46 |
| HepII•ADPHep•FDHLA (pose1) |  | 5.26 ± 0.71 |
| HepII•ADPHep•FDHLA (pose2) |  | 6.17 ± 0.32 |

Figure S1: **Sequence alignment of *E. coli* HepI and *E. coli* HepII from crystal structures.** Pairwise sequence alignment of *E. coli* HepI (PDB: 6DFE) and HepII (PDB: 1PSW) with residues in red boxes denoting identical residues.

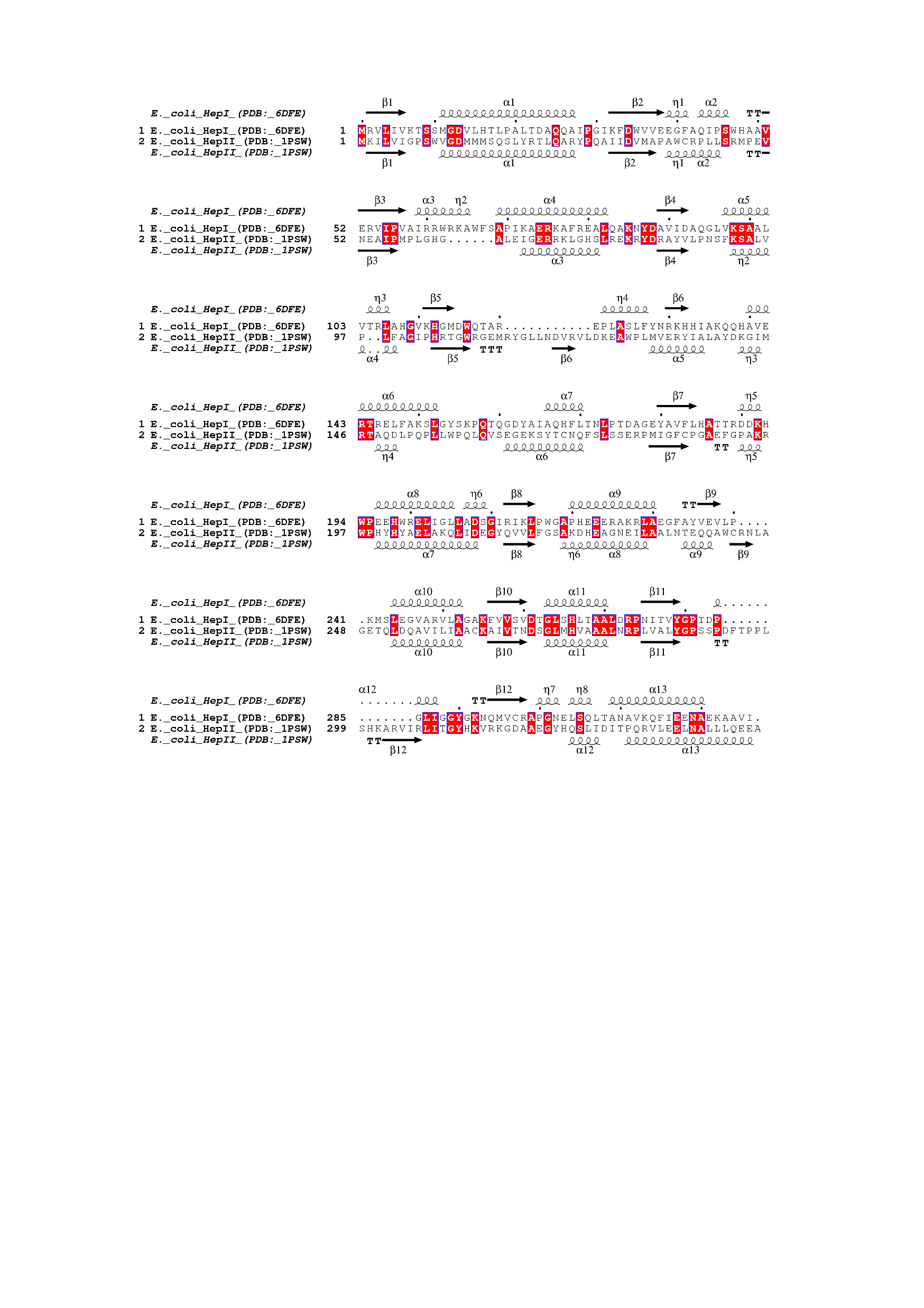

Figure S2**: Representative sequences of HepI multiple sequence alignment.** First 5 sequences of multiple sequence alignment for HepI with the query sequence (first sequence) from the crystalized *E. coli* HepI (PDB: 6DFE) and the secondary structural elements of the first sequence are represented immediately above the sequence. Residues in red boxes denote strictly conserved residues, and ones in bolded/boxed are greater than 70 similar.

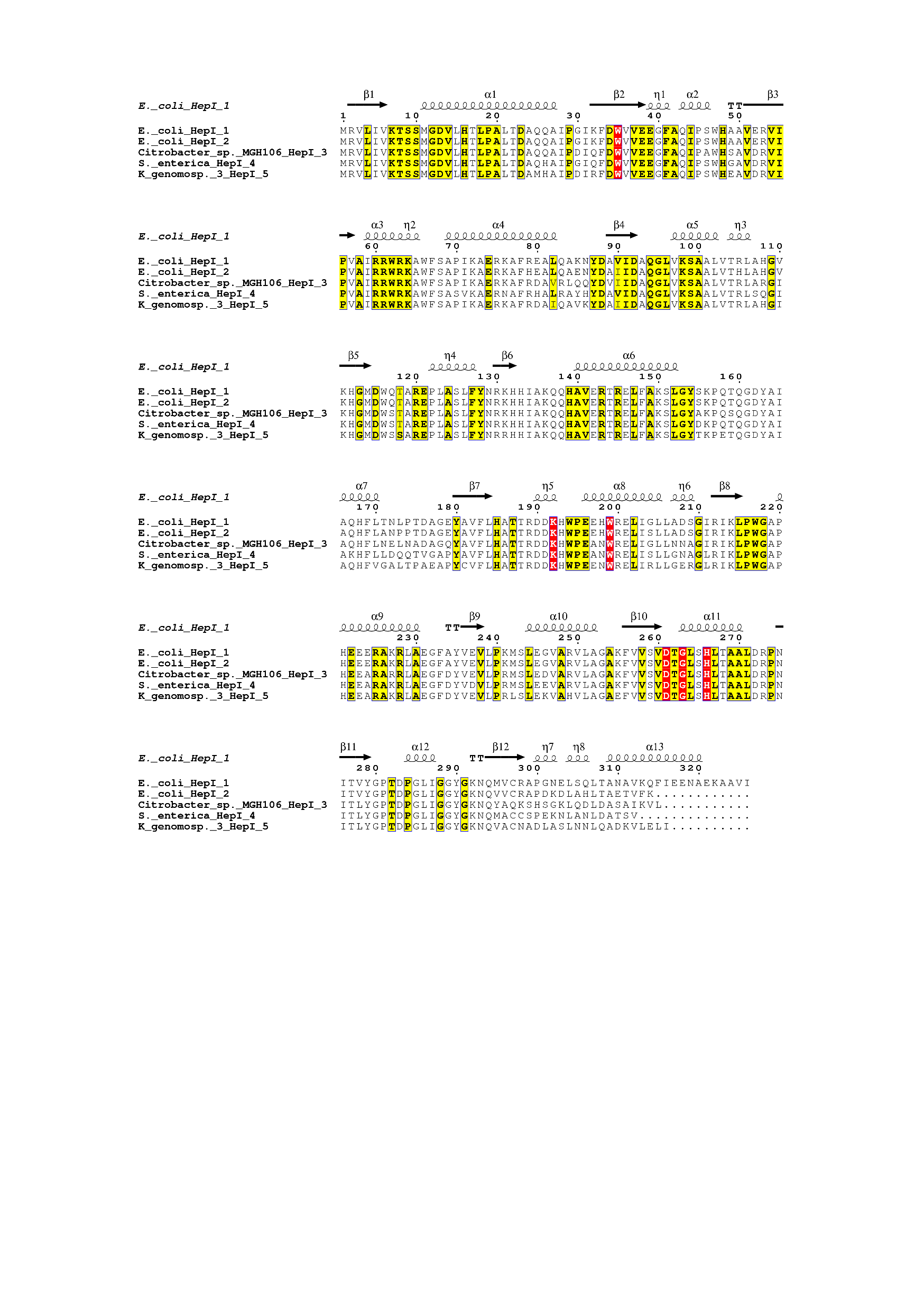

Figure S3: **SDS-PAGE gel of HepII purification.** Representative purification of E. coli HepII with fractions from flowthrough (FT), washes (W1,W2,W3), and elution (E1) separated on a sodium dodecyl sulphate polyacrylamide gel electrophoresis (SDS-PAGE) to confirm successful isolation of pure protein as demonstrated in the last lane (E1).

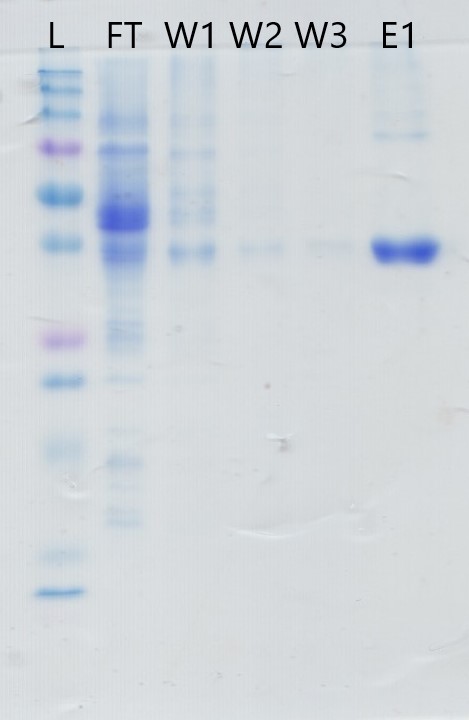

| Figure S4: **Circular dichroism spectra of HepII at varying temperatures with substrates/products.** HepII secondary structure profile from 0-95° C for **(A**) apo, **(B)** ODHLA, **(C)** ADP-Hep, **(D)** ADP, **(E)** ODH_2_LA. | |
| --- | --- |
| **A)** | 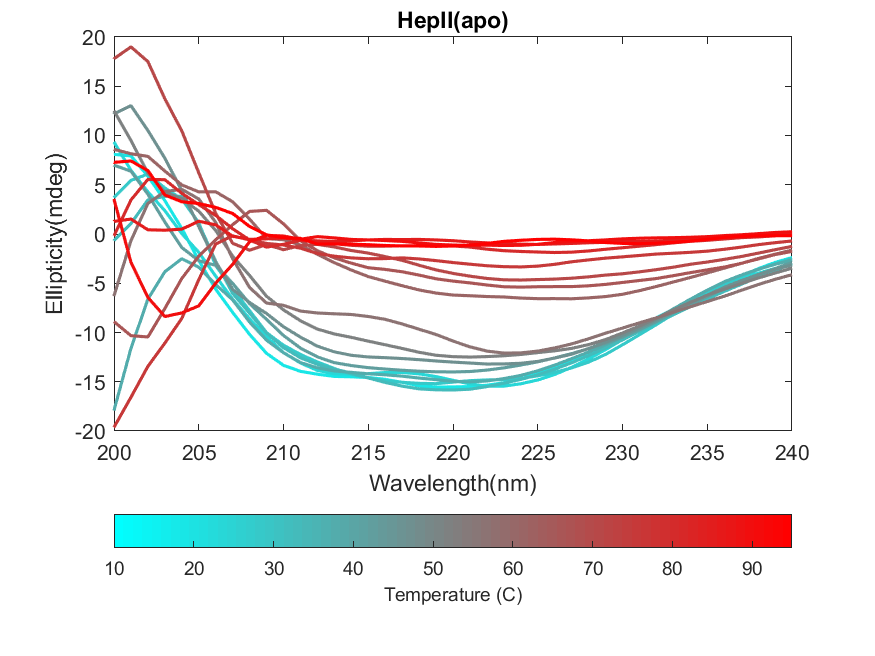 |
| **B)** | 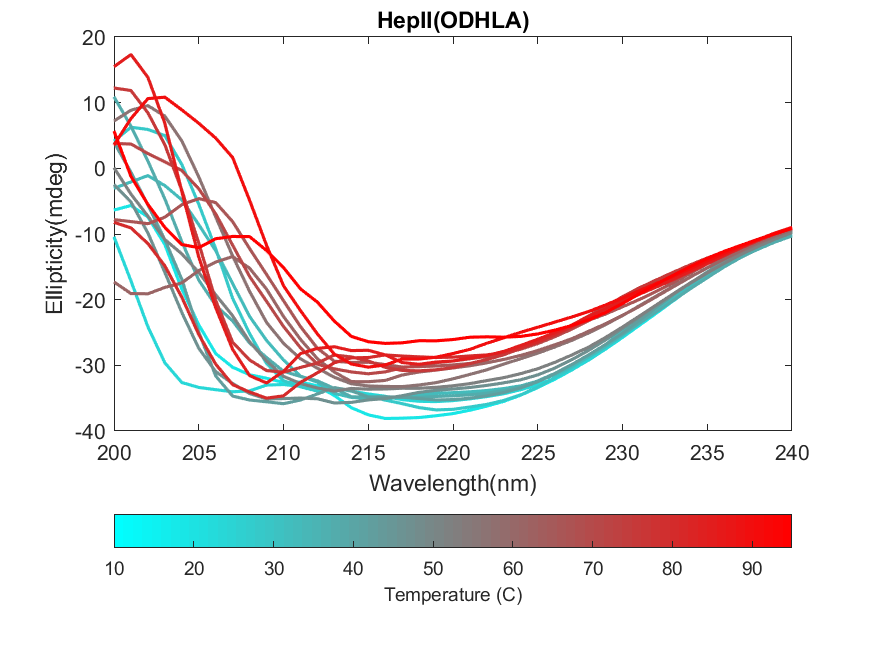 |
| **C)** | 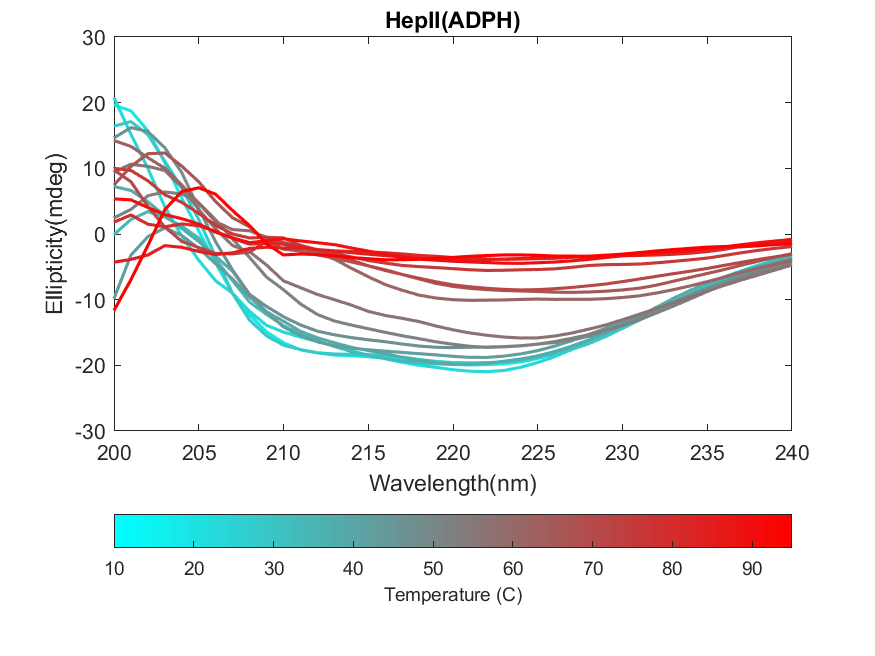 |
| **D)** | 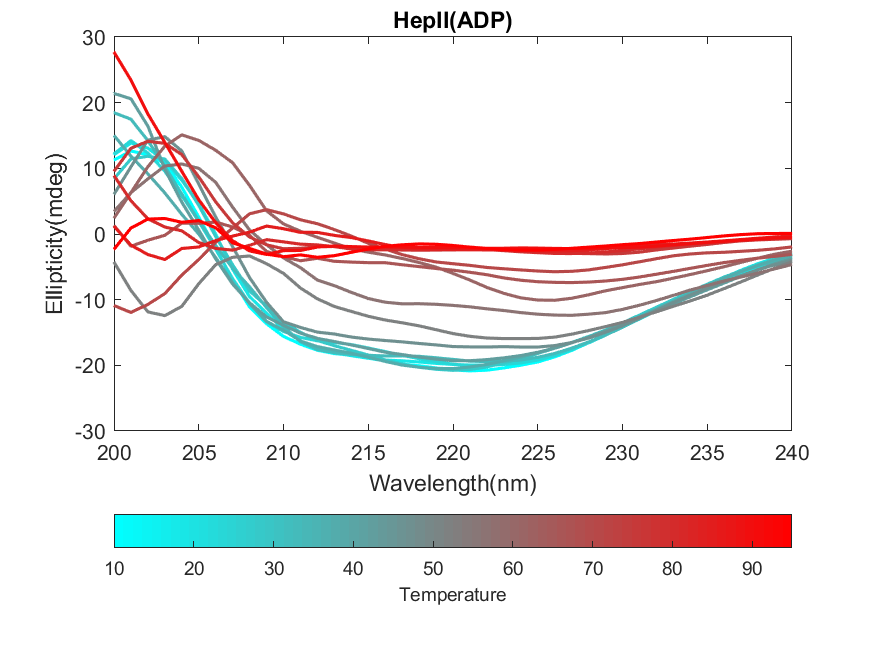 |
| **F)** | 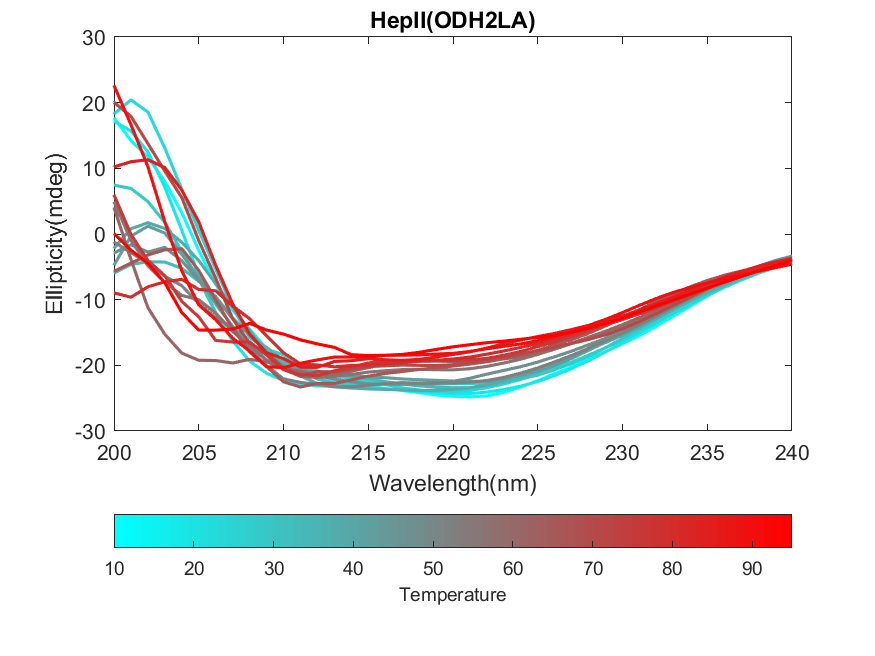 |

Figure S5: **ODHLA-HepII binding curve.** Binding curve of O-deacylated Hep-Kdo2-Lipid A (ODHLA) to HepII as determined by quenching of tryptophan fluorescence as a function of ODHLA concentration.

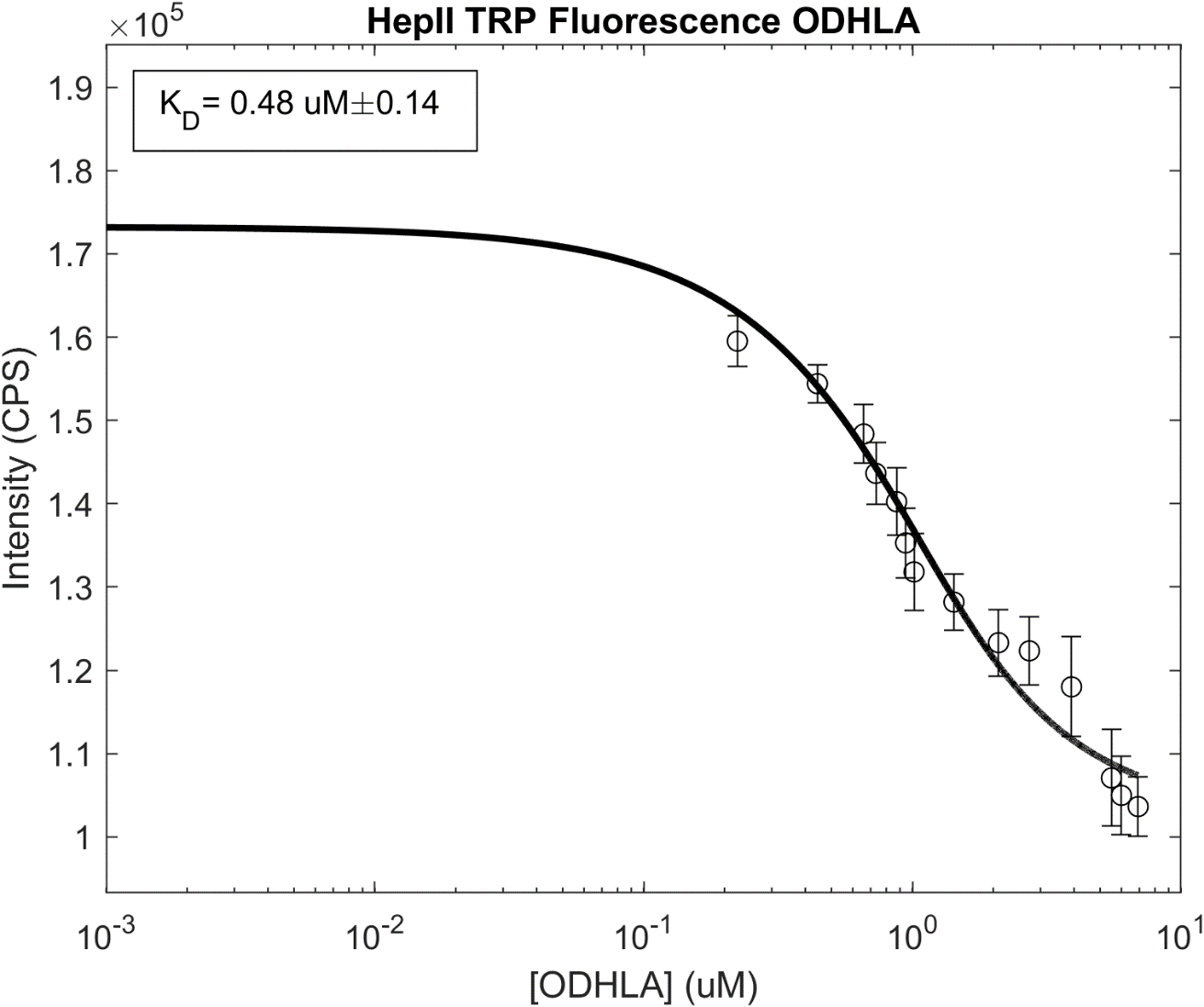

| Figure S6: **Reaction rate profiles HepII and acceptors/donors.** Michaelis-Menten curve for HepII varying fully acylated HLA with (w/ ADP-Hep: **A)**, w/ADP-Man: **B)**) and without triton x 100 (w/ ADP-Hep: **C)**, w/ADP-Man: **D)**), O-deacylated HLA (w/ ADP-Hep: **E)**, w/ADP-Man: **F)**) and O-deacylated HLA with varying **(G)** ADP-Hep and **(H)** ADP-Man . | |
| --- | --- |
| **A)** | 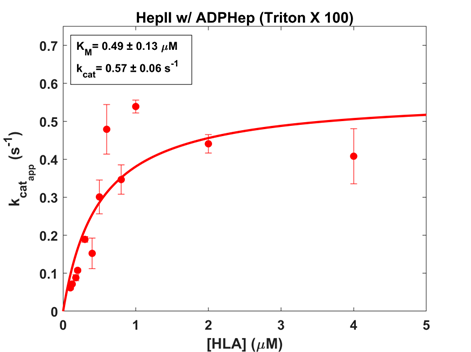 |
| **B)** | 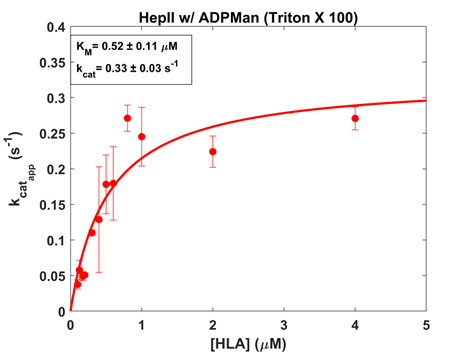 |
| **C)** | 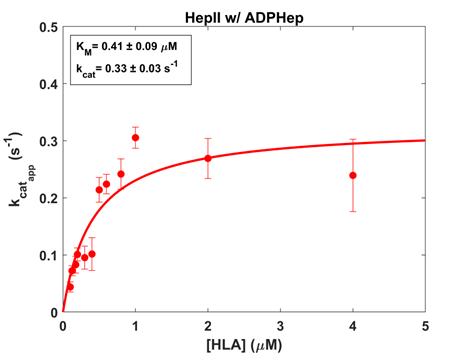 |
| **D)** | 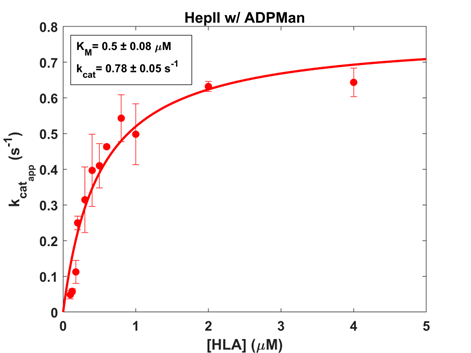 |
| **E)** | 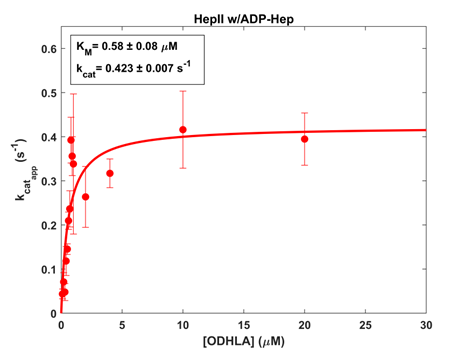 |
| **F)** | 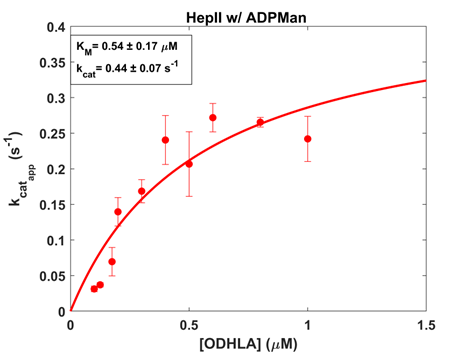 |
| **G)** | 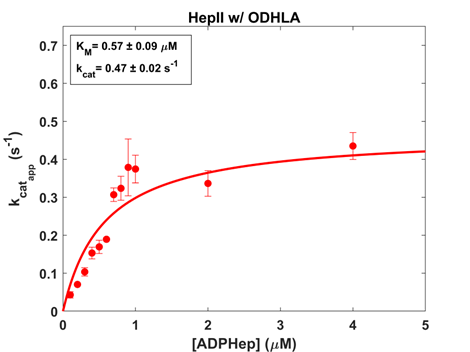 |
| **H)** | 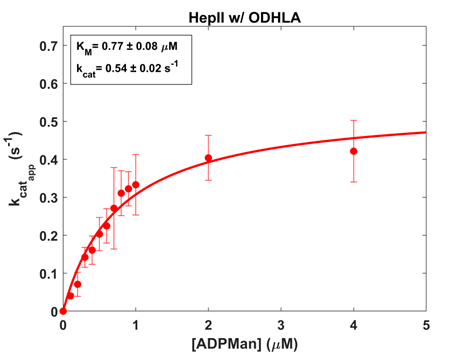 |

Figure S7: **Mass spectrometric characterization of ODHLA (acceptor, substrate).** ESI mass spectra of O-deacylated Hep-Kdo_2_-Lipid A.

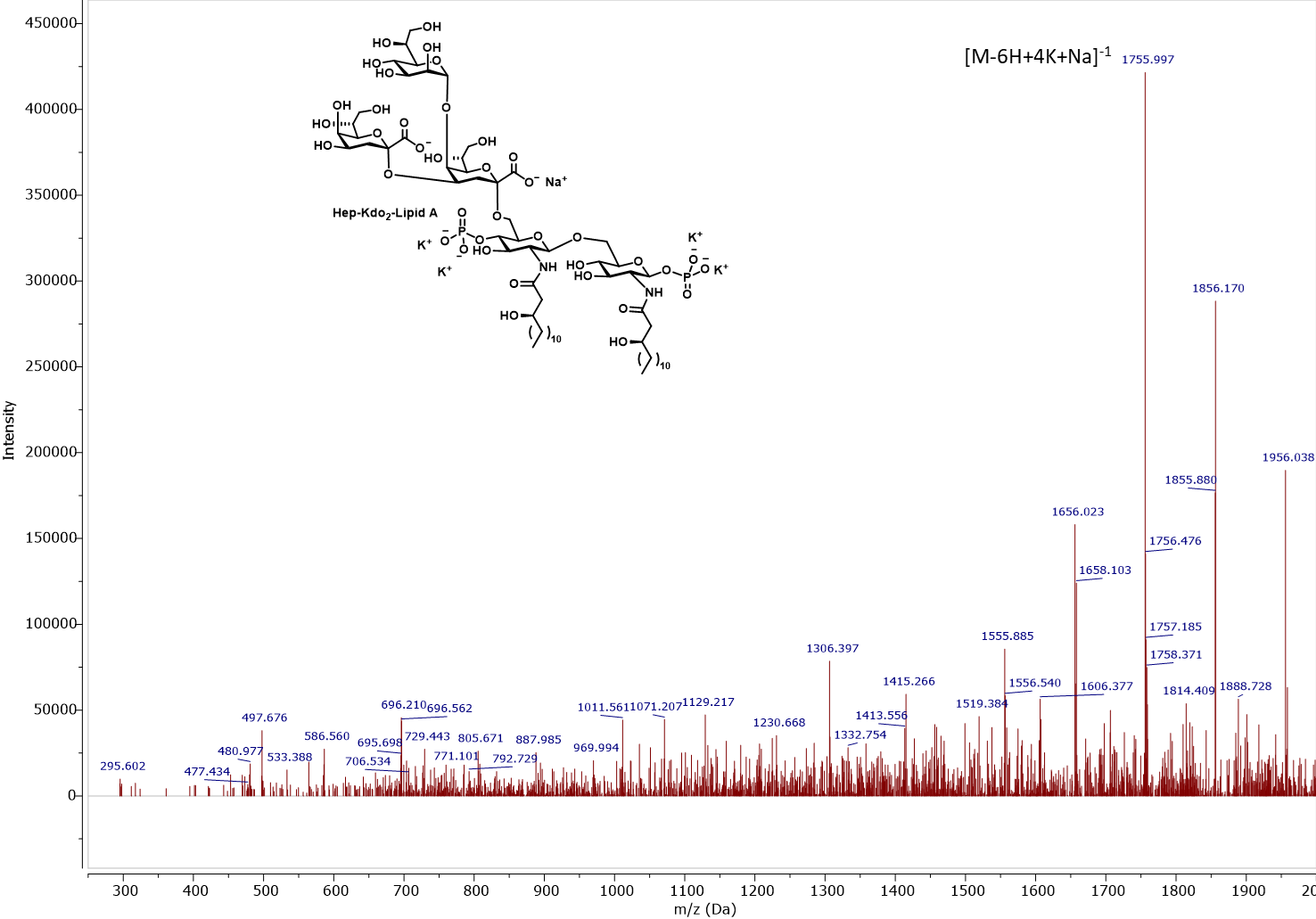

Figure S8: **Mass spectrometric characterization of ODH_2_LA (acceptor, product)**. ESI mass spectra of O-deacylated Hep2-Kdo2-Lipid A.

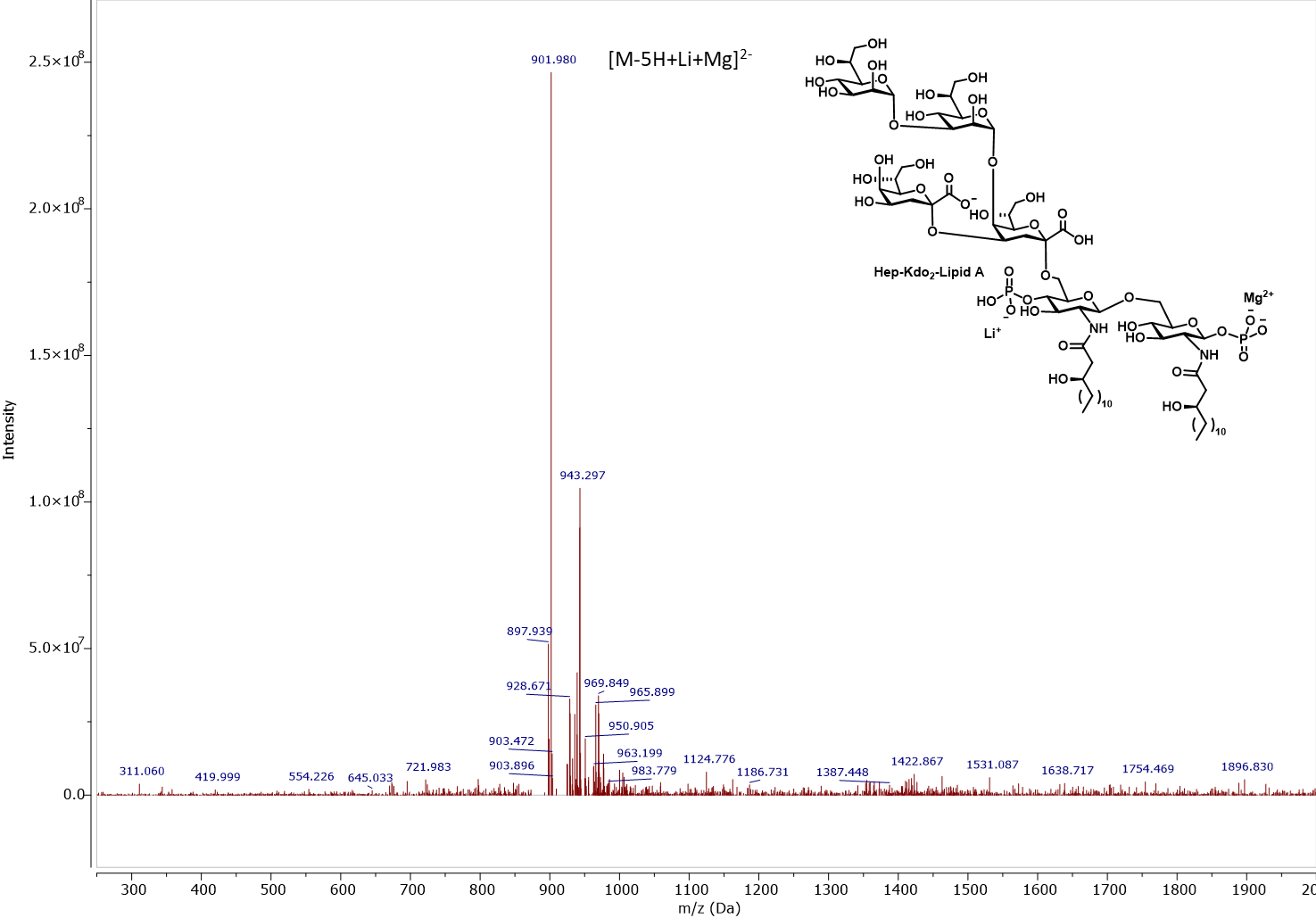

Figure S9: **Time course analysis of ADP-Man biosynthesis.** ^31^P NMR spectral analysis enabled monitoring ADP-Man biosynthesis and phosphatase digestion to degrade excess ATP, ADP, and AMP.

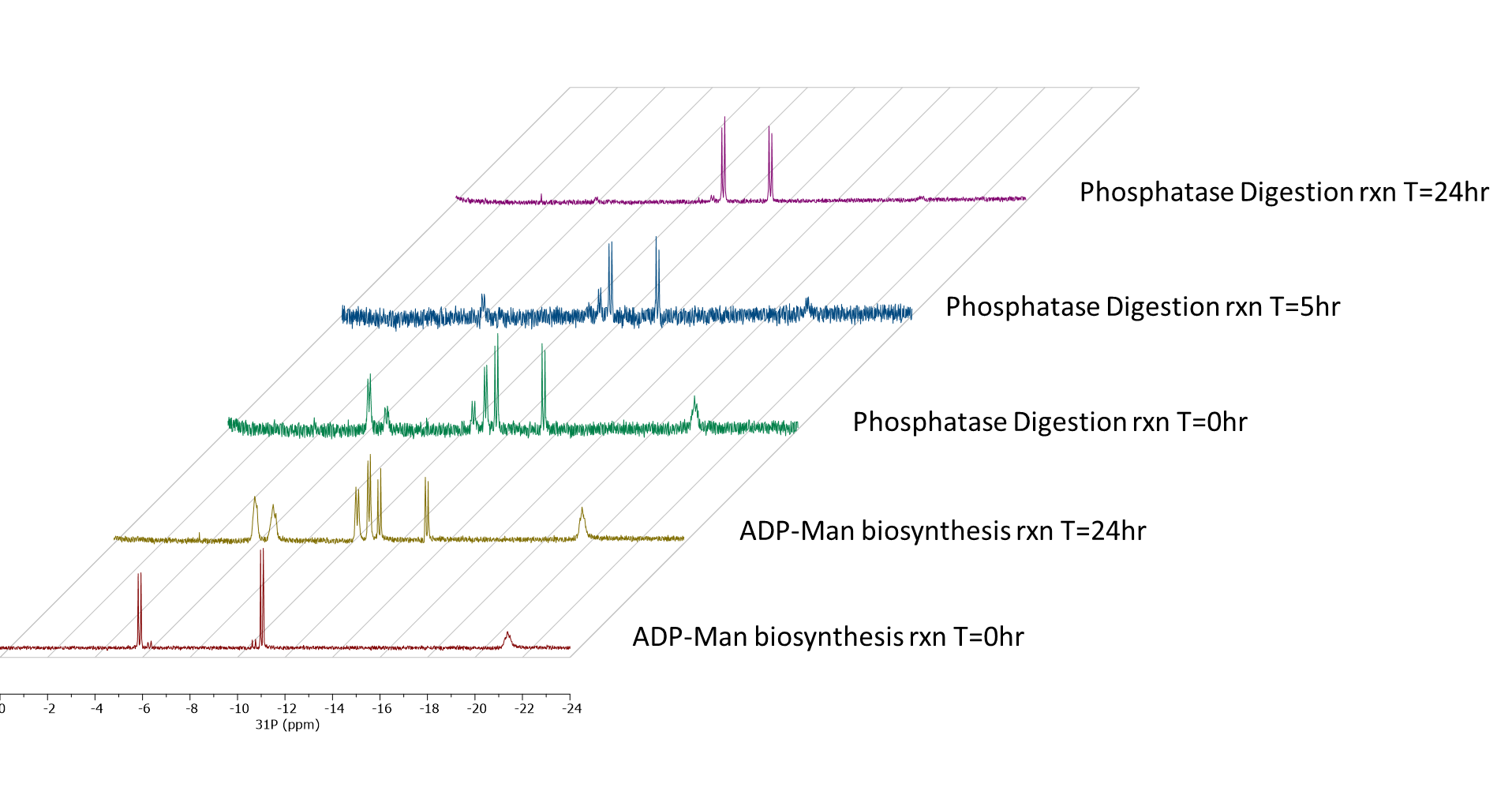

Figure S10: **Mass spectra of ADP-Man (donor, substrate).** ESI mass spectra of ADP-Man.

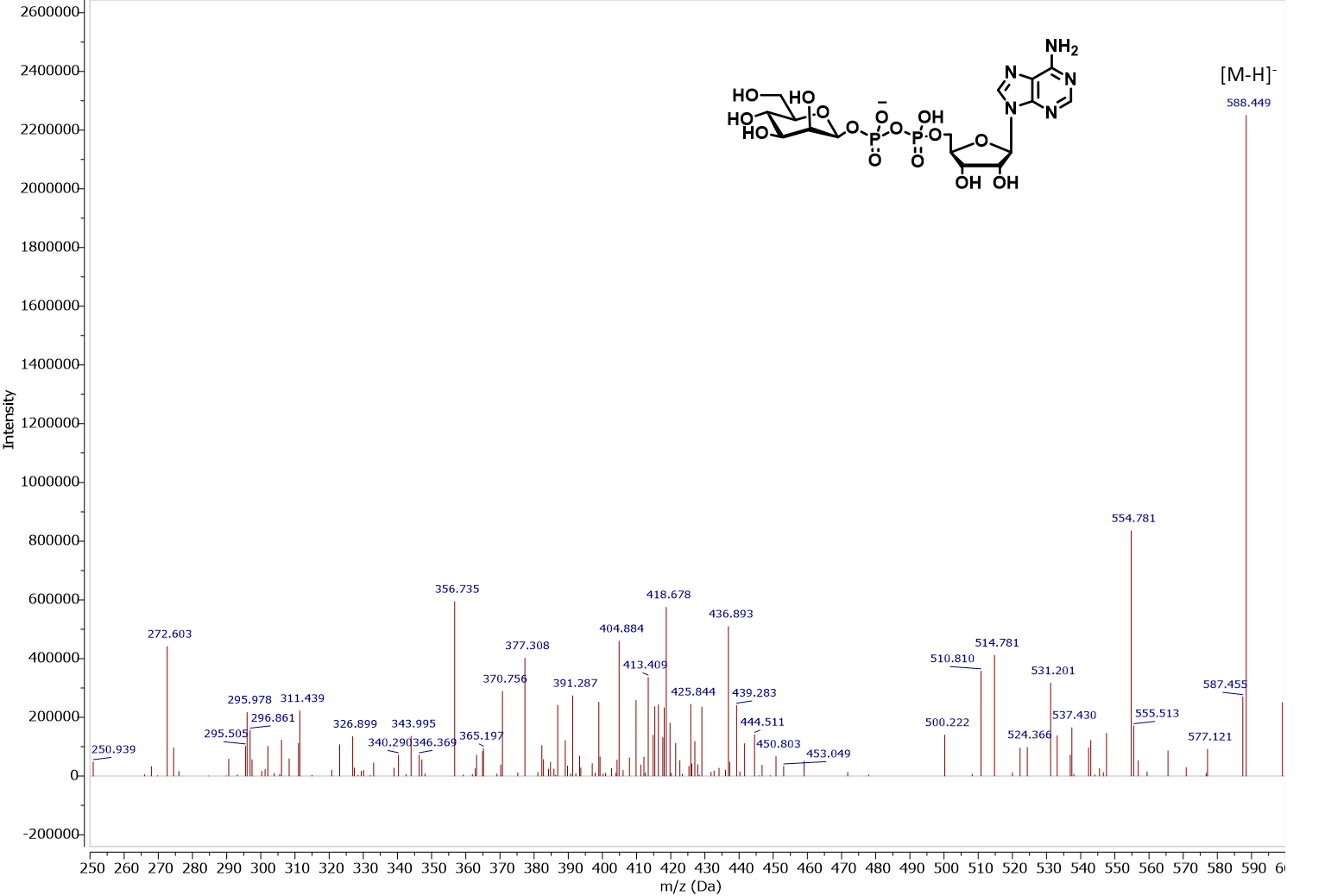

| Figure S11: **RMSD and RMSF of HepII trajectory.**  **(A)** Backbone RMSD and **(B)** C^α^ RMSF of HepII apo and both poses of the HepII•ADP-Hep•FDHLA complex. | |
| --- | --- |
| **A)** | 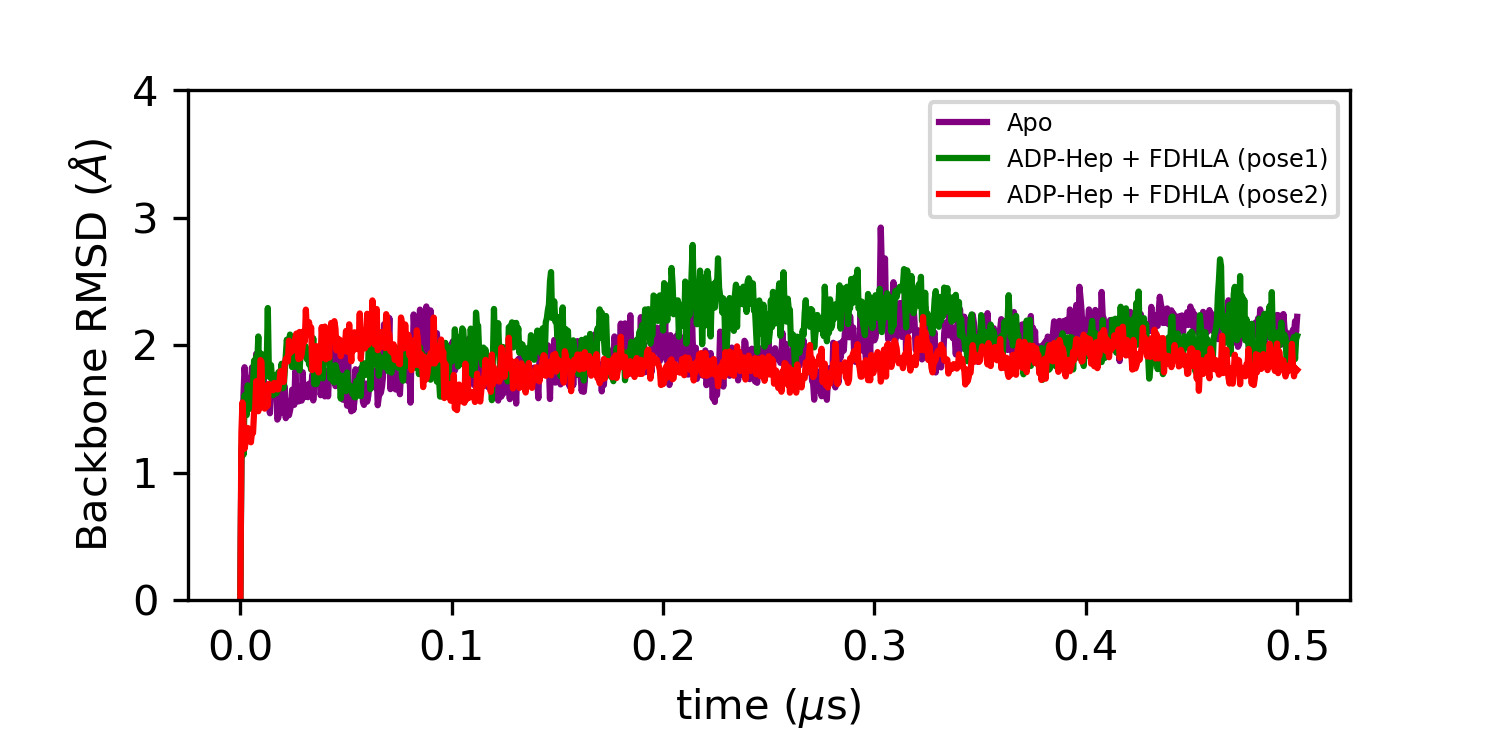 |
| **B)** | 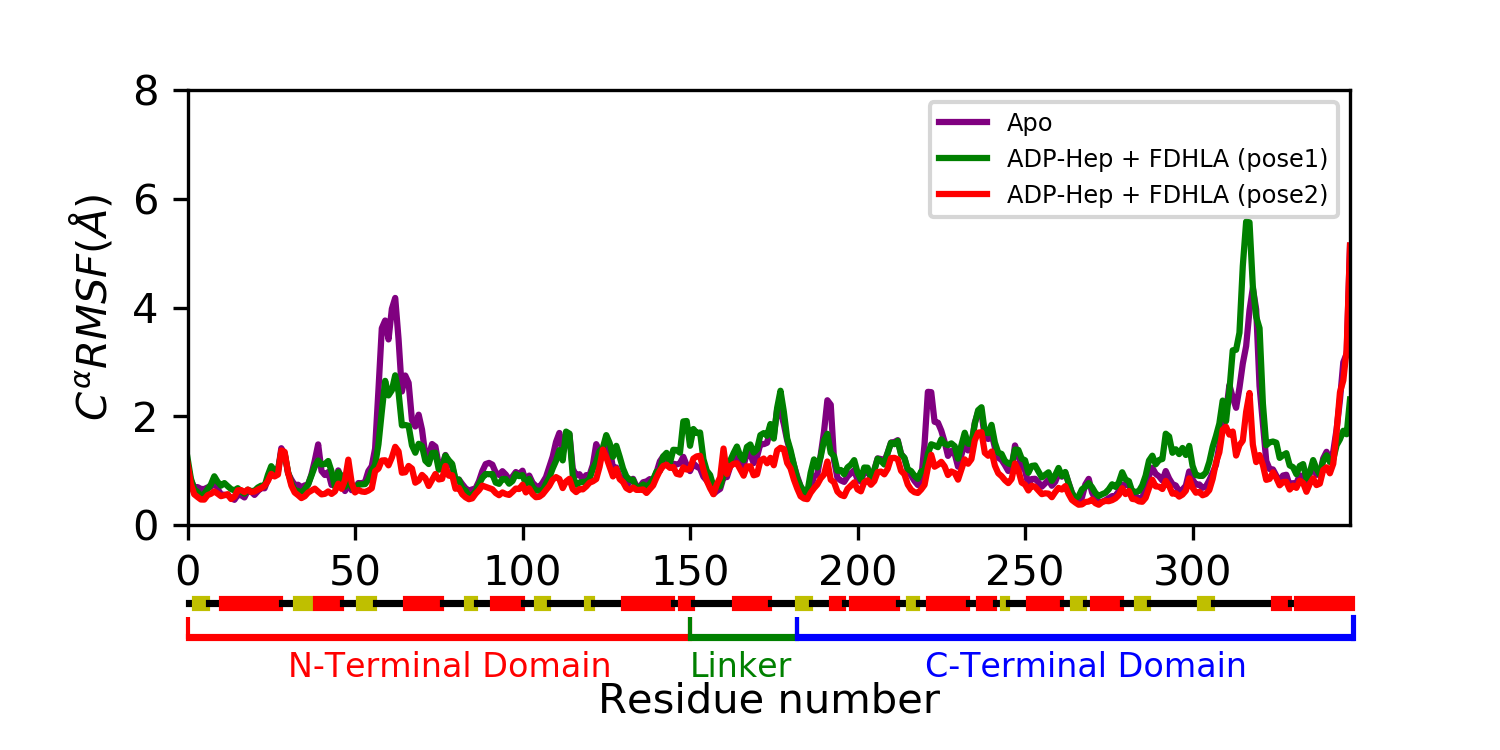 |

| Figure S12: **Protein/ligand (pose2) interactions from simulation.** Interactions between HepII sidechains and **(A)** FDHLA (pose2) or **(B)** ADP-Hep over the course of the trajectory. | |
| --- | --- |
| **A)** | 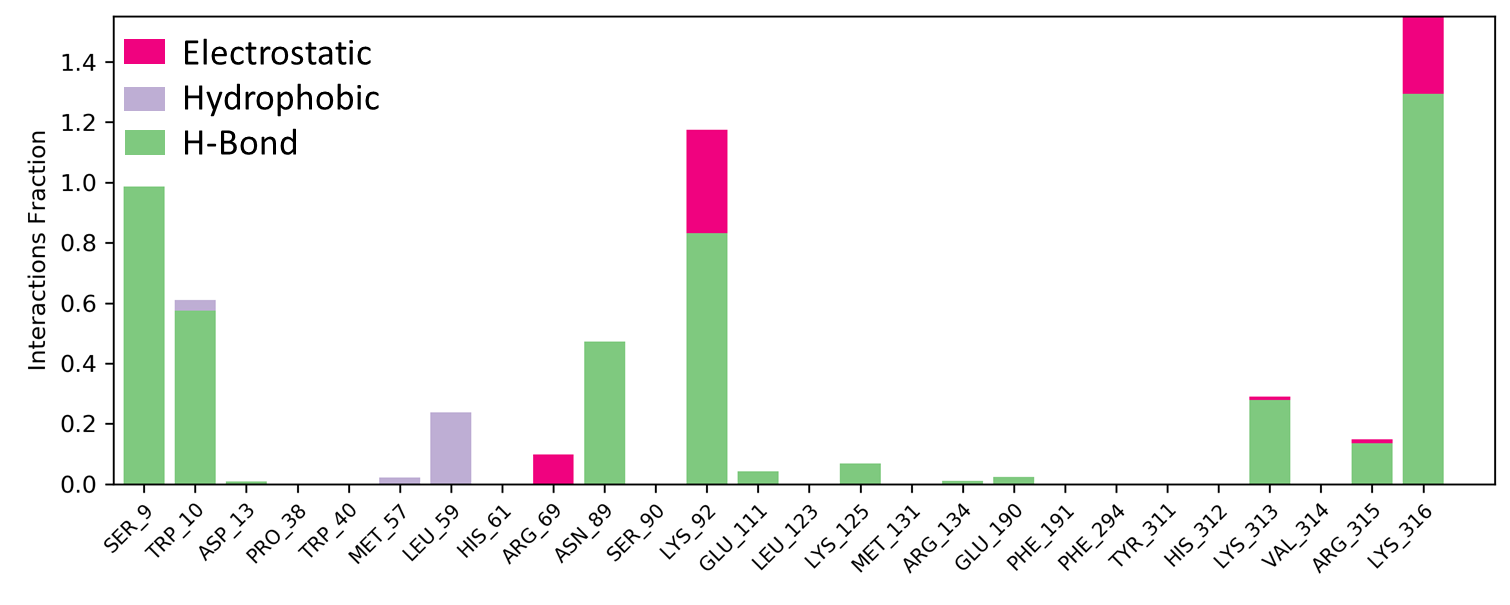 |
| **B)** | 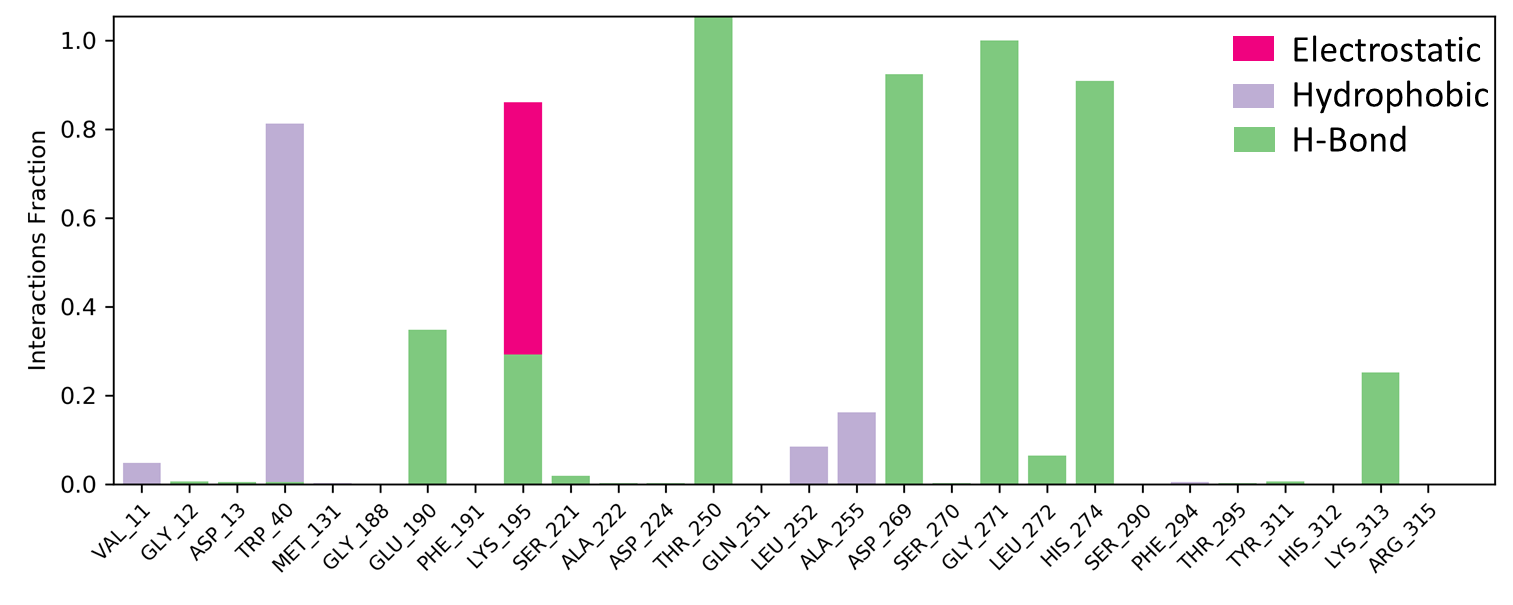 |

| Figure S13: **Ligand interaction diagram of substrates (pose1) to HepII sidechains across MD trajectory.** Contacts between HepII sidechains and **(A)** FDHLA (pose1) or **(B)** ADP-Hep. | |
| --- | --- |
| **A)** | 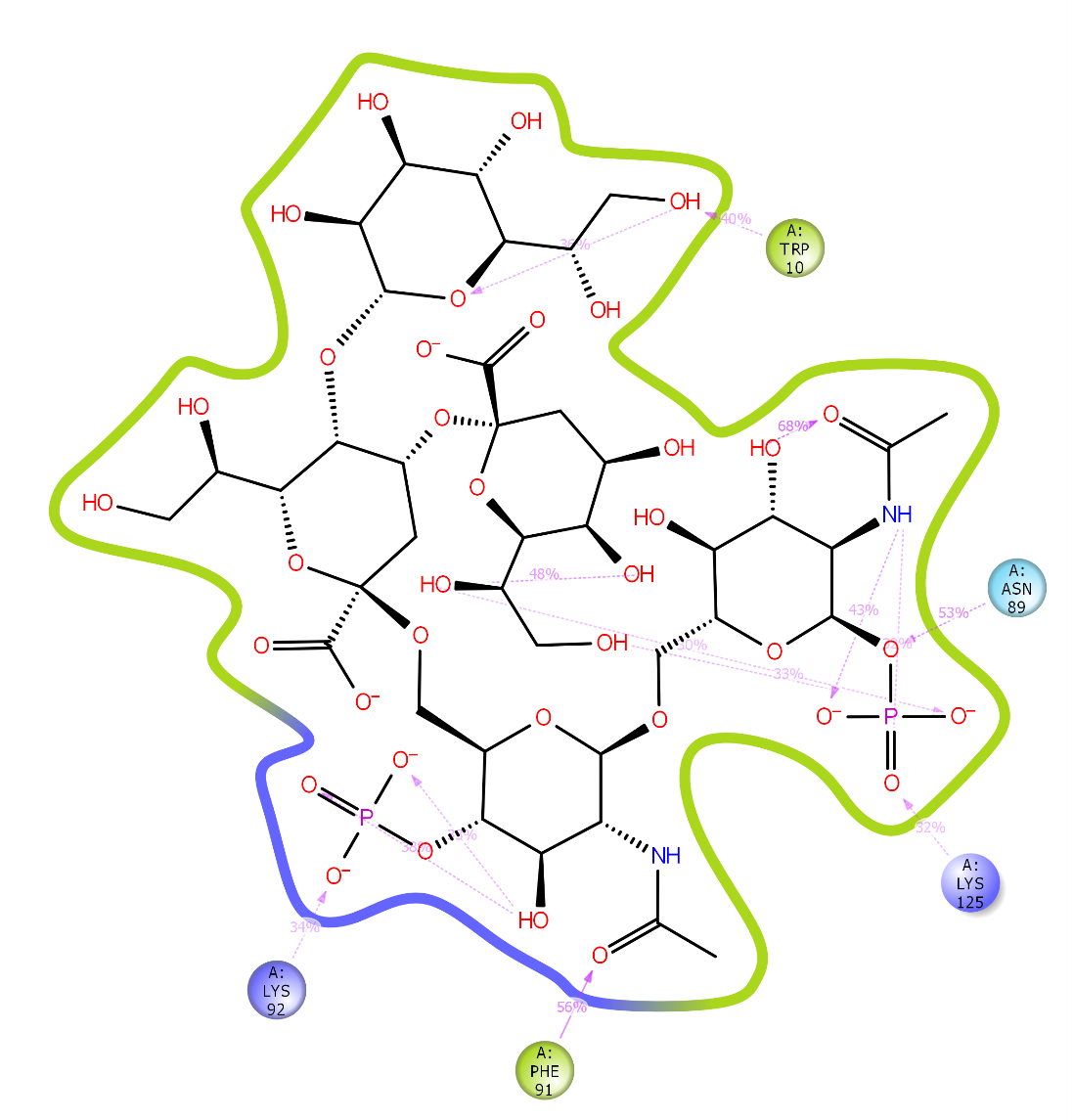 |
| **B)** | 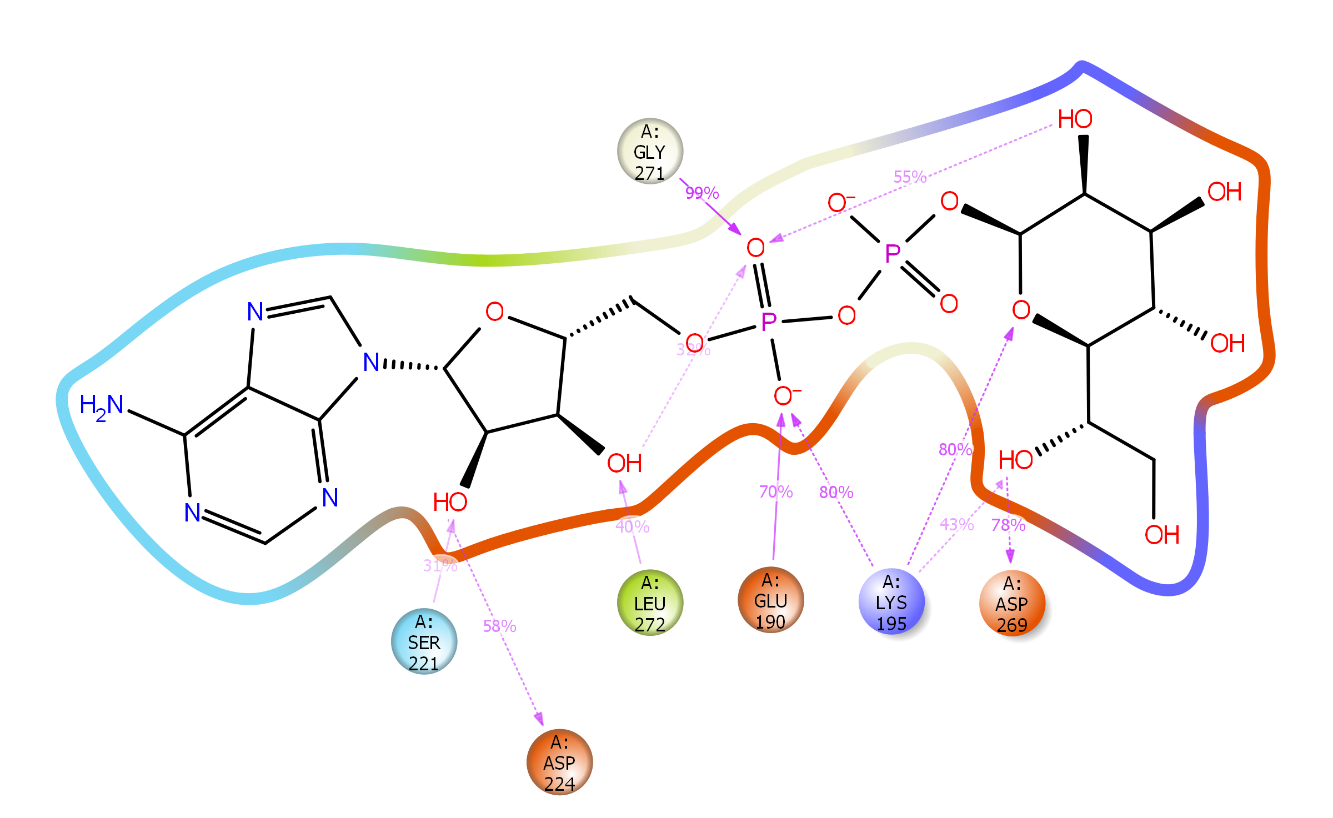 |

| Figure S14: **Protein/ligand (pose1) interactions from simulation.** Interactions between HepII sidechains and **(A)** FDHLA (pose1) or **(B)** ADP-Hep over the course of the trajectory. | |
| --- | --- |
| **A)** | 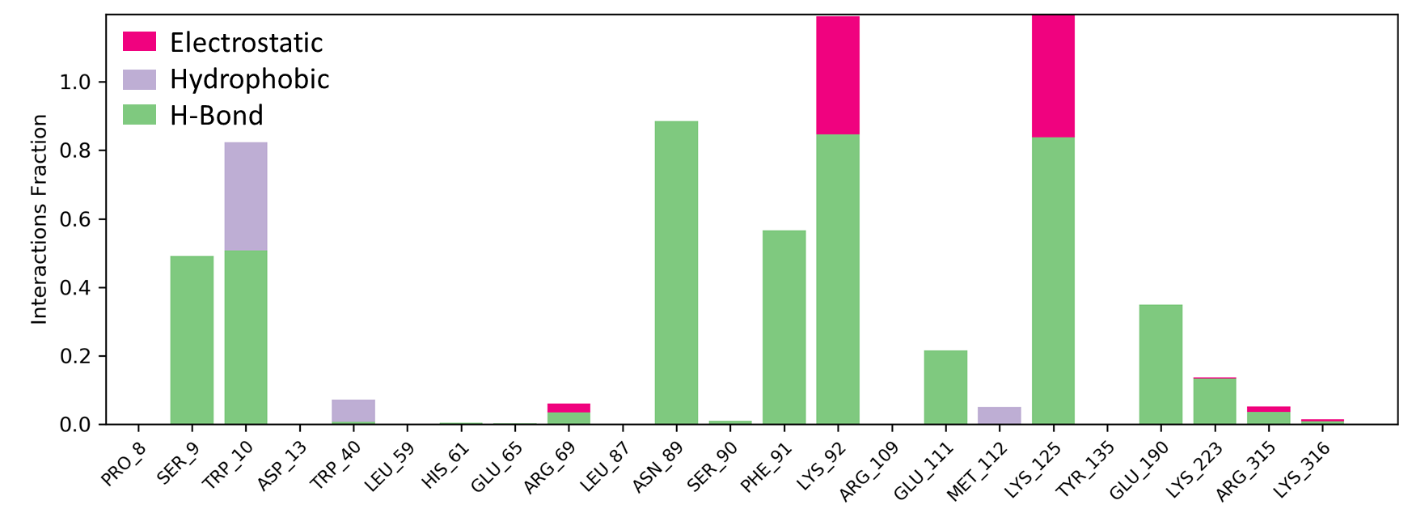 |
| **B)** | 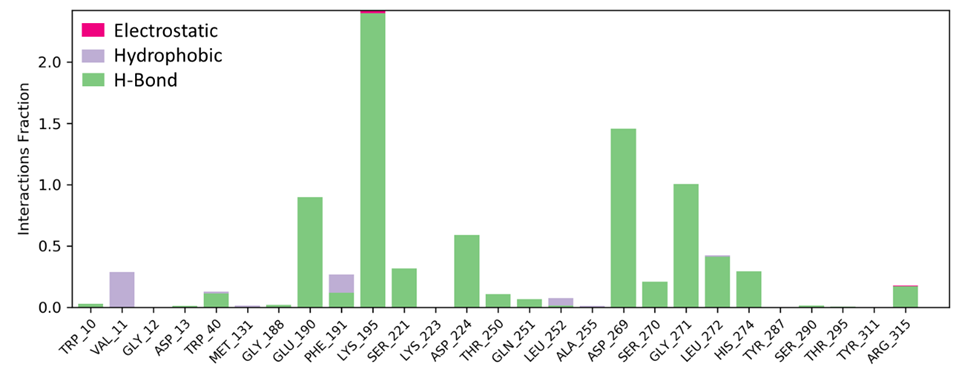 |

| Figure S15: **Secondary structural changes observed in MD simulations of apo and ligand bound complexes.** Structure of HepII (A) apo and (B) ternary complex representative frame demonstrating an increase in α helicity in the 60s region (black box) due to ligand induced stabilization of this highly dynamic loop. | | | |
| --- | --- | --- | --- |
| **A)** | 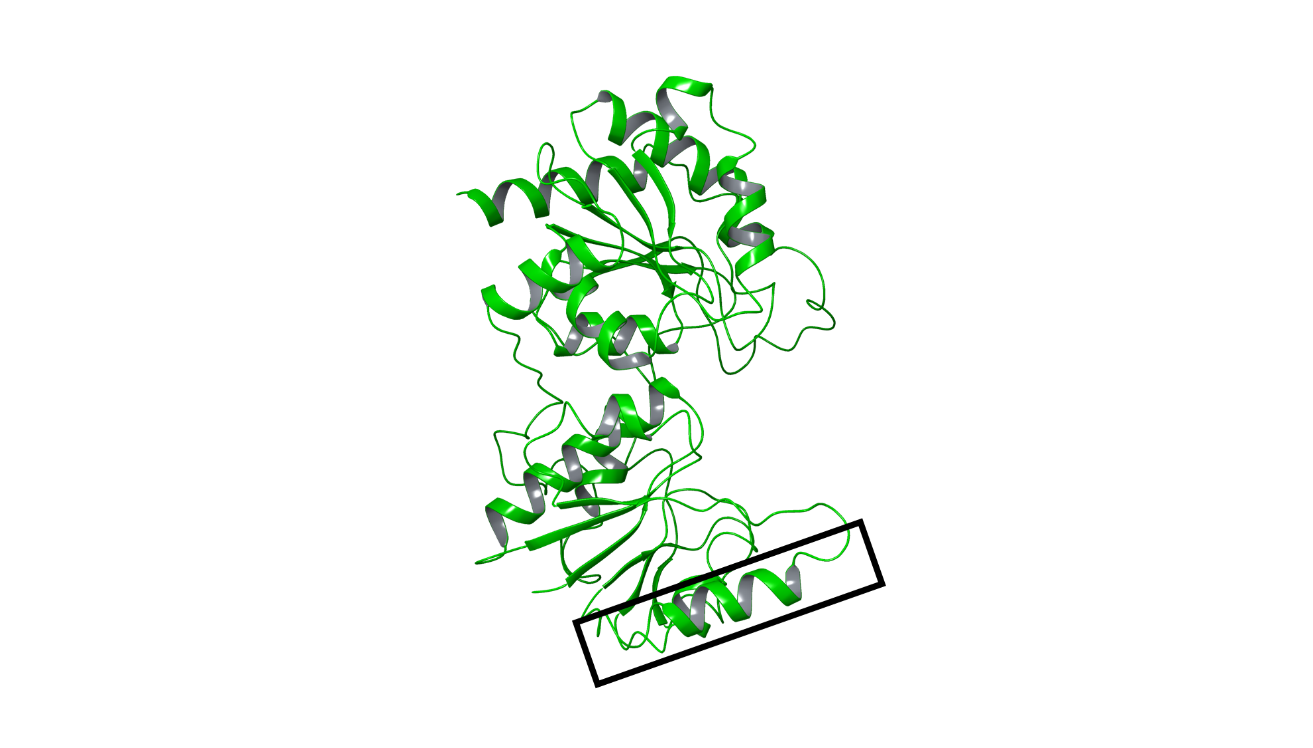 | **B)** | 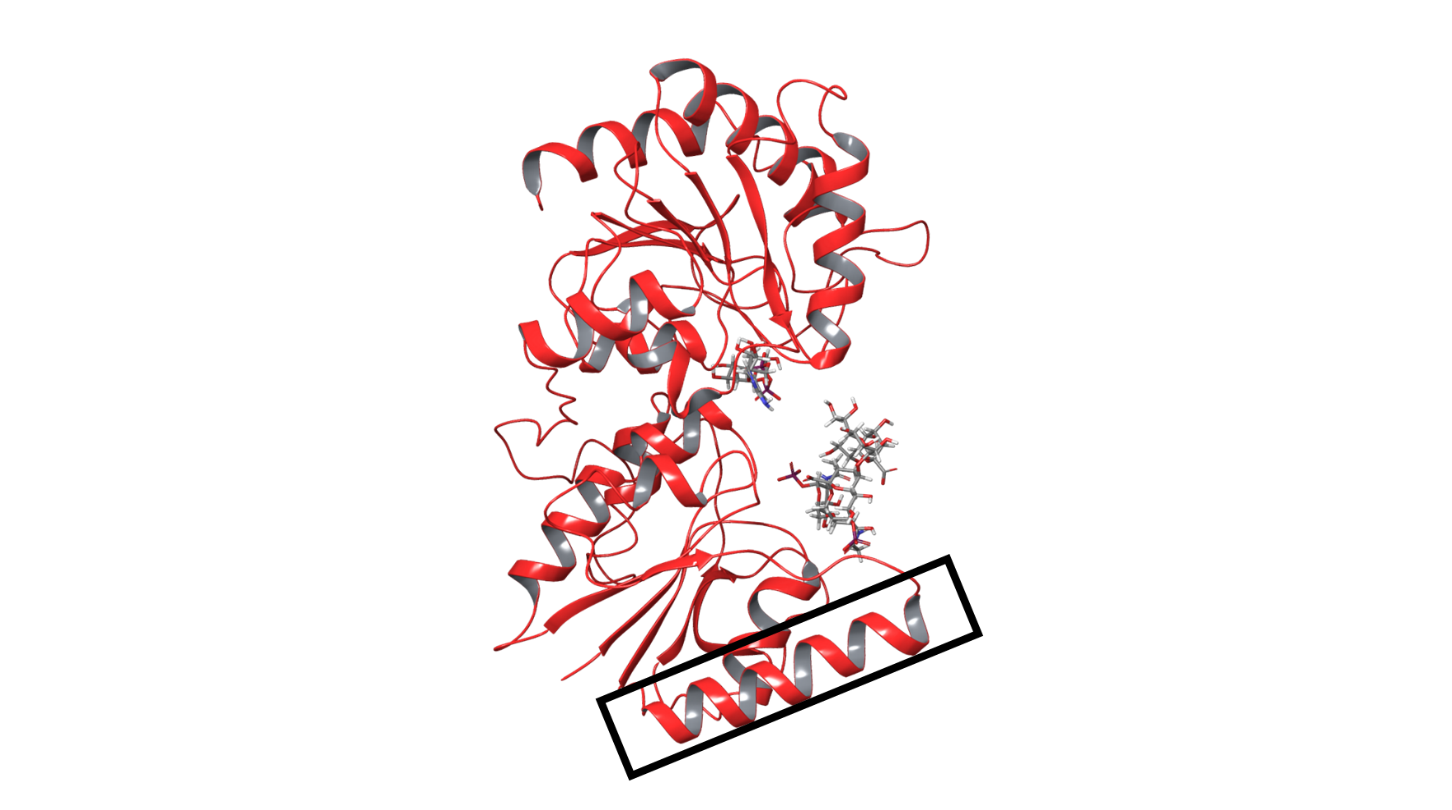 |
